# A statistical model for quantitative analysis of single-molecule footprinting data

**DOI:** 10.1101/2025.05.20.655044

**Authors:** Evgeniy A. Ozonov, Laura Gaspa-Toneu, Antoine H.F.M. Peters

## Abstract

The binding of sequence-specific TFs (TF) to genomic DNA is fundamental to gene regulation. Emerging single-molecule footprinting (SMF) technologies such as the NOMe-seq and Fiber-seq assays offer unique opportunities for acquiring quantitative information about binding states of TFs and nucleosomes at single-DNA-molecule resolution. Contrasting bulk epigenomic profiling methodologies, SMF enables better molecular characterization of inherently stochastic processes of protein-DNA interactions. Despite the many advantages that SMF technologies bring for studying mechanisms of gene regulation, rigorous statistical models for the analysis of datasets generated using these technologies are still missing. Here, we introduce a novel statistical framework designed for inference of footprint lengths and predictions of footprint positions for unbiased quantitative analysis and interpretation of SMF datasets. We carried out comprehensive computational simulations of SMF experiments and identified experimental parameters that are critical to footprint detection. Finally, we demonstrate the power of this statistical approach for the analysis of genome-wide and amplicon-based NOMe-seq datasets generated for mouse embryonic stem cells.

## INTRODUCTION

The regulation of transcription plays crucial roles in maintaining cellular homeostasis (1) and ensuring the appropriate timing, location, and levels of gene activities during organismal development and differentiation processes (2). This regulation is primarily mediated by transcription factors (TFs) - DNA-binding proteins that recognize specific motifs in the genome and modulate transcription. In eukaryotes, the binding of TFs to their cognate sites occurs in the context of local chromatin states influenced by DNA methylation (3–5), post-translational modifications (PTMs) of histones (6–9), and nucleosome positioning (10–12).

In the past decades, genome-wide high-throughput assays such as ChIP-seq (13, 14), MNase-seq (15, 16), ATAC-seq (17), and others (18) have provided remarkable insights in the locations of TFs and nucleosomes in various cell types (19). Nonetheless, these assays have relatively low genomic resolution (hundreds of bps) and only measure average enrichments across larger cell populations. Hence, they are unable to characterize important regulatory aspects by TFs such as the inherent stochasticity of binding events within cell populations and the co-occupancy of TFs and nucleosomes on single chromatin molecules. In contrast, emerging Single-Molecule Footprinting (SMF) methods provide high-resolution insights into the location of DNA-binding proteins on individual chromatin fibers and can achieve nearly single-base resolution (20–28).

Typically, SMF assays involve two key steps (Figure 1A). The first step involves enzymatic marking of positions on DNA fibers that are unprotected by DNA-binding proteins, such as histones or TFs. For example, the original study by Kelly et al. (20), introducing the Nucleosome Occupancy and Methylome Sequencing assay (NOMe-seq), utilized a bacterial GpC methyltransferase (M.CviPI). This enzyme adds selectively methyl groups to accessible cytosines within GpC dinucleotides, enabling footprint profiling on single chromatin fibers in eukaryotes. Since endogenous 5mC marks at GpC sites are rare in eukaryotic genomes, unlike 5mC at CpG sites, any detected 5mC methylation at GCH sites (where H represents a non-guanine nucleotide, thus excluding GpCpG sites that may undergo endogenous CpG methylation) can be confidently attributed to M.CviPI activity. This ensures that the identified methylation marks reflect regions of chromatin devoid of DNA-binding proteins.

**Figure 1:**
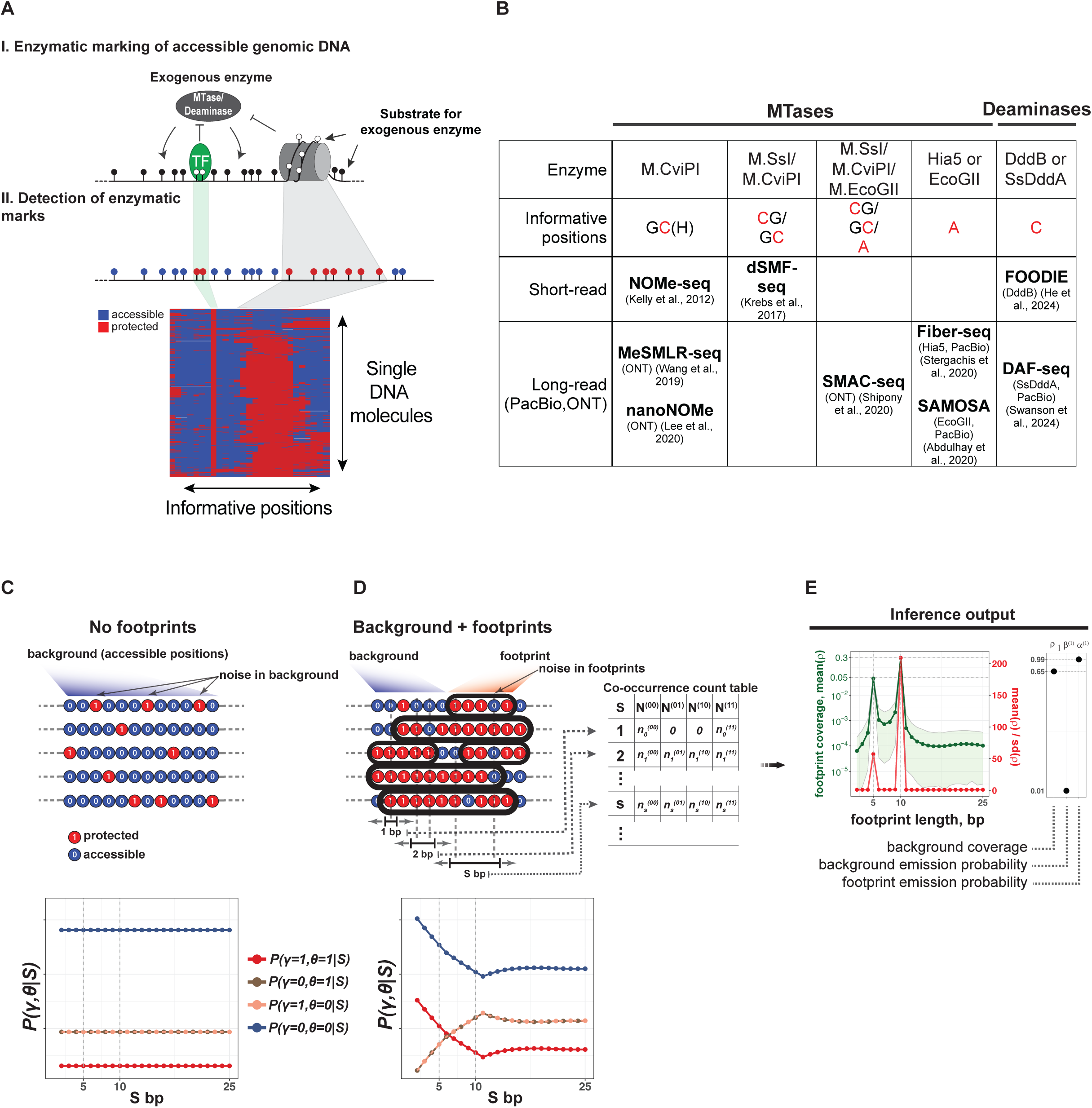
Statistical inference of footprint lengths from SMF datasets. (A) Schematic description of SMF assays. Substrate nucleotides for enzymatic marking, e.g. cytosines in GpC contexts for M.CviPI, adenines for M.EcoGII/Hia5 and cytosines for DddB/SSDddA, are represented as lollipops colored in black for unprotected and white for protected positions. The heatmap with raw data illustrates a reduced representation of SMF data where only informative positions are shown and non-informative positions are omitted. Colors of the heatmap depict protection (red) or accessibility (blue) at informative positions. (B) The table represents an overview of currently existing SMF assays with the first row describing enzymes used for marking accessible positions and the second row representing corresponding substrate nucleotides with informative positions highlighted in red. The third and fourth rows list assays that use short-read or long-read sequencing technologies for detecting modified sequences. The usability of a particular assay for SMF experiment depends on endogenous DNA methylation, e.g. dSMF-seq is suitable only for organisms lacking endogenous m5CpG methylation (21). (C) An *in-silico* example of SMF data which consists of only noisy background without footprints. All positions are accessible but due to supposed technical noise we observe a low percentage of protected (unmarked) positions. In this case, joint probabilities 𝑃(𝛾, 𝜃|𝑆) of observing different combinations of protected and unprotected positions at distance S bp are constant and do not depend on S, assuming statistical independence between positions. Joint probabilities 𝑃(𝛾 = 0, 𝜃 = 1|𝑆) and 𝑃(𝛾 = 1, 𝜃 = 0|𝑆) are equal due to symmetry. (D) An *in-silico* example of SMF data consisting of noisy background and footprints with lengths of 5 bp and 10 bp. Statistical independence between positions is broken due to presence of footprints in the data, therefore probabilities 𝑃(𝛾, 𝜃|𝑆) for data with footprints depend on distance between positions 𝑆𝑆. Input data for footprint spectral analysis is a co-occurrence count table which represents frequencies of combinations of protected and accessible positions at various distances observed in a SMF dataset. (E) An example of an output from the footprint spectral analysis for *in-silico* generated data in (D) which consists of footprints with lengths of 5 bp and 10 bp. Statistical inference is performed by Hamiltonian Monte Carlo sampling from posterior distribution or by sampling from approximating mean-field distribution obtained by variational Bayesian methods implemented in the statistical programming language Stan. Green and red curves illustrate two alternative representations of the footprint spectrum. The green curve represents mean values with 2.5%-97.5% percentiles and the red curve represents ratios between means and standard deviations across samples from the posterior distribution (or variational approximation) for footprint coverages.

The second step involves detecting the chemical modification on DNA introduced by the exogenous enzyme at accessible positions. For example, the original NOMe-seq assay (20) employed bisulfite treatment which converts unmethylated cytosines to uracils. This was followed by short-read high-throughput sequencing, enabling the differentiation of accessible and protected cytosines at GCH sites, as well as the detection of endogenous DNA methylation at WCG sites (where W represents A or T) within single chromatin fibers.

Although the NOMe-seq assay provides single-molecule information, its genomic resolution is limited to GCH sites, due to the sequence specificity of M.CviPI and the presence of endogenous CpG methylation in eukaryotes. Additionally, the use of short-read sequencing restricts the ability to assess long-range co-occupancy of DNA-binding proteins. To address these limitations, subsequent studies have explored alternative exogenous enzymes (25–28) or combinations of enzymes (21, 24), along with advanced methods for detecting enzymatic marks over longer genomic ranges within the same chromatin fibers (22–26, 28) (Figure 1B). Moreover, SMF assays have been adapted for integration into single-cell multi-omics studies (29–31).

Despite substantial advancements in improving genomic resolution and extending molecule length, the computational analysis of SMF datasets remains a significant challenge. The limited genomic resolution of the selected enzyme, incomplete enzymatic marking of accessible positions, and errors in their detection contribute to high noise levels in SMF datasets (32). To address these issues, recent approaches have incorporated statistical models such as Hidden Markov Models (33, 34), methods that analyze average accessibility across defined genomic windows (21, 32), and neural network-based techniques for more robust identification of enzymatic marks (34, 35). However, these methods are limited in their ability to estimate footprint lengths present in SMF data directly, to distinguish between nucleosome and TF footprints and often rely on prior knowledge of TF binding sites.

Here, we introduce a novel statistical framework designed to identify footprints and their locations in single chromatin molecules in an unbiased and quantitative manner. The first component of our model is focused on what we term “footprint spectral analysis”. This involves identifying the footprints present in the dataset and estimating the noise levels within accessible (background) and inaccessible (footprints) regions. By addressing this, our method provides insights into the abundance of different footprint types and the variability in the data.

Building on this foundation, the second component of our approach determines the footprint positions within individual molecules in the dataset. Given the known lengths, abundances, and noise levels derived from the first component, the second part of the model assigns probabilities to each footprint for covering (or starting from) every position within a molecule, effectively disentangling footprints of different lengths and accurately mapping their spatial organization. Together, these two components enable a quantitative analysis of SMF datasets, offering opportunities for high-level characterization, detailed positional insights into footprinting patterns, and molecule classification to estimate binding rates of nucleosomes and TFs.

Furthermore, we developed two R packages: *fetchNOMe* for the fast retrieval of NOMe-seq data from BAM files, and *nomeR* for the analysis of SMF datasets using the statistical approach described in this study.

## RESULTS

### A statistical model for inference of footprint lengths in SMF datasets

The first component of our statistical framework is the footprint spectral analysis that is aimed at estimating how much of a given SMF dataset is composed of free unprotected DNA (background) and possible footprints regardless of their positions within molecules, and optionally to estimate noise levels within background and footprints observed in the given SMF dataset. In other words, we wish to estimate a “footprint spectrum” which represents coverages for footprints with lengths within defined ranges.

To achieve this, we assume that the observed single-molecule footprinting (SMF) data are generated by a random process in which footprints of different lengths are placed along an infinitely long DNA sequence according to their underlying abundances, which we aim to estimate. Each footprint then produces a pattern of protected or accessible states at each position it covers, based on specific emission probabilities (see Supplementary Text for details, Figure S1).

The underlying principle of inferring footprint spectra in SMF datasets is based on the derivation of joint probabilities 𝑃𝑃(𝛾, 𝜃|𝑆) of observing a pair 𝛾, 𝜃 ∈ {0,1} representing a combination of accessible (encoded with 0) or protected (encoded with 1) positions at distance 𝑆 bp from each other for the described stochastic process (Supplementary Text, Figure S3).

Indeed, when we consider an SMF dataset with noisy background and containing no footprints, then joint probabilities 𝑃(𝛾, 𝜃|𝑆) is constant for any distance 𝑆 bp, assuming statistical independence between positions (Figure 1C).

In contrast, footprints introduce statistical dependency between positions and the probability 𝑃(𝛾, 𝜃|𝑆) becomes a function of the distance between positions 𝑆, footprint abundances and noise levels (Figure 1D).

Therefore, using the derived joint probability 𝑃(𝛾, 𝜃|𝑆) for the stochastic process described above (see eq. 36 in Supplementary Text) we can construct a Bayesian model (Figure S6) and use observed frequencies 𝑁^(𝛾θ)^ at various distances 𝑆 (Figure 1D) for inferring footprint spectra and optionally emission probabilities for background and footprints (Figure 1E).

### Sensitivity of statistical footprint spectral analysis to experimental factors in SMF assays

Protocols for SMF assays typically involve multiple steps, each introducing experimental noise into the resulting dataset. For instance, incomplete or excessive enzymatic marking, failures in detecting enzymatic marks, and sequencing errors contribute to noise in both accessible and protected positions. Additionally, the density of informative positions depending on the selected enzyme (e.g., methyltransferases M.CviPI, Hia5, M.EcoGII, or deaminases DddB and SsDddA; Figure 1B) and the sequence composition of regions of interest can be limiting. Finally, the sequencing depth is influenced by the choice of sequencing platform and the available experimental budget. Consequently, the quality of resulting SMF datasets critically depend on these experimental factors, which directly impact the ability to perform statistical analyses.

To evaluate the performance of our model for footprint spectral analysis and assess its sensitivity to these experimental factors, we conducted *in-silico* simulations for two experimental scenarios (Figure 2A). These scenarios involved footprints corresponding to nucleosomes and TFs, which are common DNA-binding protein complexes investigated in SMF assays. Both scenarios included a long (150 bp) footprint, representing nucleosomes, that lacked positional preference and covered 50% of positions within a region of interest (Figure 2A). In the second scenario, a short (50 bp) footprint, representing TFs, covered 5% of the positions, occupying a specific site within the region of interest and competing with the randomly positioned 150 bp footprint (“anchored 50 bp & random 150 bp footprints” in Figure 2A).

**Figure 2:**
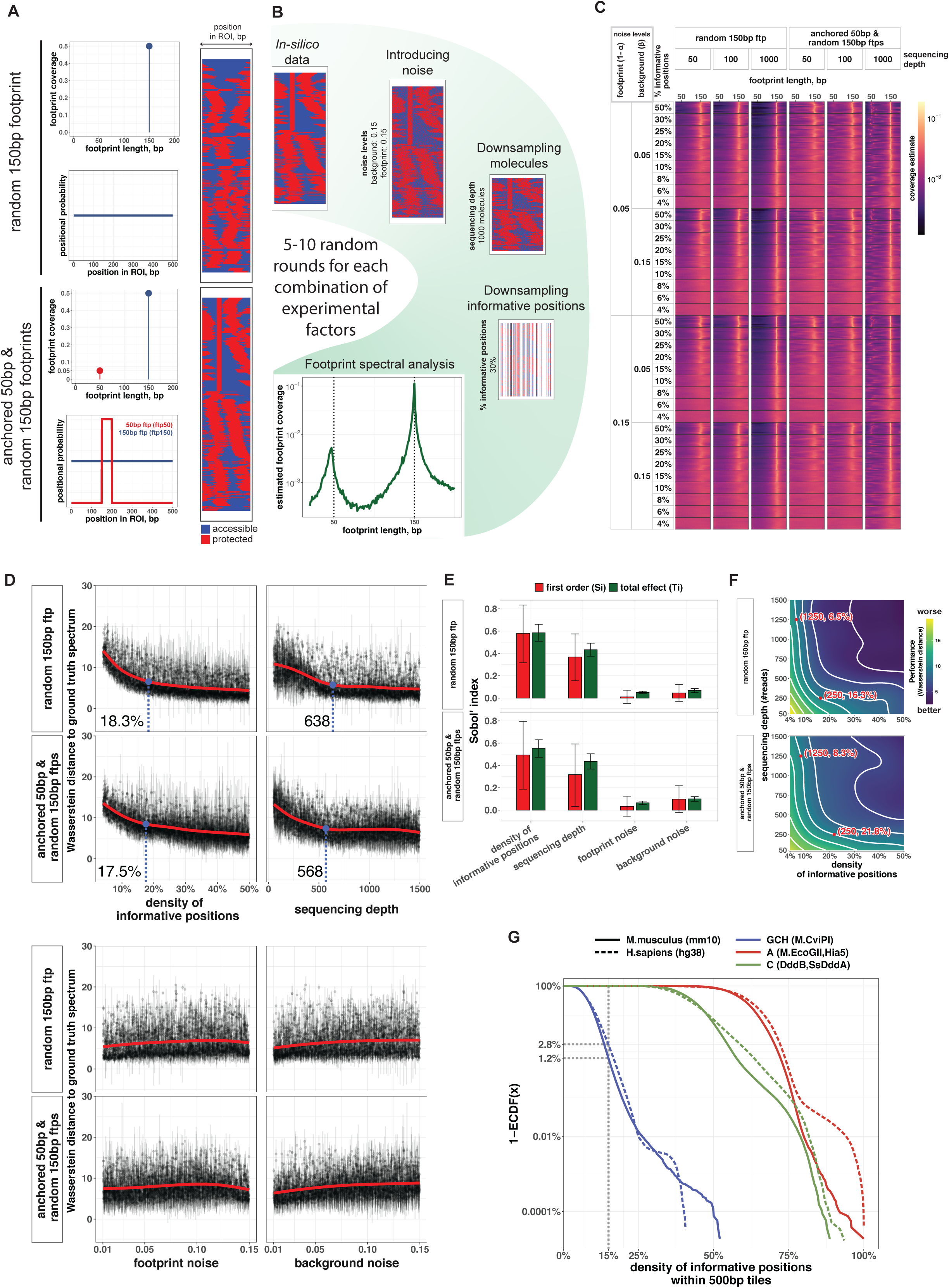
Sensitivity of footprint spectral analysis to experimental factors. (A) Generation of *in-silico* SMF datasets for two experimental scenarios. The first scenario involves a footprint of length 150bp (“nucleosome”) that covers 50% of positions and is randomly placed within a 500 bp region of interest (ROI). The second scenario, in addition to a 150bp footprint, also includes a footprint of length 50bp (“TF”) that covers 5% of positions and can occupy only its binding site at a particular position within the ROI. *In-silico* SMF data are generated by randomly placing footprints according to footprint spectra (panels for footprint coverage) and positional probability profiles along ROI (see Methods for details). At this stage of *in-silico* SMF data generation it is assumed that all positions within the ROI are informative, lacking noise within background and footprints. (B) Introducing experimental biases and errors into SMF datasets generated in (A). We independently introduce the following experimental factors into generated SMF datasets: i) we randomly mutate values within background and footprint positions according to emission probabilities (noise levels), ii) we randomly downsample the number of sequenced reads, and iii) we randomly downsample informative positions within ROI according to selected density of informative positions (see Methods). For each combination of experimental parameters, we generate 5 to 10 random *in-silico* SMF datasets. (C) Heatmaps illustrating footprint spectra inferred from *in-silico* SMF datasets generated using the procedure described in (B) for different sequencing depths, densities of informative positions and emission probabilities for background and footprints. We generated 10 random *in-silico* SMF datasets for every combination of experimental parameters using the procedure described in (B) and each row within heatmaps corresponds to a footprint spectrum for a particular *in-silico* SMF dataset. (D) Scatters illustrating performance of the footprint spectral analysis at recovering “ground truth” spectra for the two scenarios in (A). Number of experimental parameter sets that included various combinations of densities of informative positions, sequencing depths, emission probabilities for background and footprints were randomly sampled using Sobol’ method and 5 random *in-silico* SMF datasets were generated for every combination of experimental parameters for the two scenarios in (A) using the same method illustrated in (B). Inference of footprint spectrum was done for each SMF dataset generated for a particular combination of experimental parameters and similarity to “ground truth” spectra illustrated in (A) was estimated using Wasserstein distance (lower values represent better performance). Each point and error bars show mean and standard deviation of Wasserstein distances to “ground truth” spectra across 5 random *in-silico* SMF datasets. Panels illustrate how Wasserstein distances to “ground truth” spectra depend on each experimental factor included in the simulation for the two scenarios in (B). (E) Sobol’ global sensitivity of inference algorithm performance at recovering “ground truth” footprint spectra to experimental parameters. The first order (red) and total (green) Sobol’ indices were calculated for each experimental factor based on simulations described in (D). (F) Performance of footprint spectral analysis as measured by Wasserstein distance to ground-truth spectra (lower values represent better performance) for combinations of density of informative positions (X-axis) and sequencing depth (Y-axis). Contour lines designate areas with similar Wasserstein distances for various densities of informative positions and sequencing depths. (G) Percentage of 500bp regions (Y-axis) in the mouse (solid lines, mm10) and human (dashed lines, hg38) genomes with densities of informative positions above a cutoff on the X-axis for SMF assays using MTases M.CviPI (cytosines within GCH context, in blue), M.EcoGII or Hia5 (adenines, in red) and deaminases DddB or SsDddA (cytosines, in green). Substrate nucleotide motifs for each enzyme were matched on both strands.

We began by generating random footprint configurations for 3,000 chromatin fibers based on the defined experimental scenarios (Figure 2A). Each simulated SMF dataset for a specific combination of experimental parameters was created in the following manner. First, we introduced noise by randomly mutating accessible and protected positions according to the specified levels of background and footprint noise (Figure 2B). Next, to simulate limiting experimental factors, such as sequencing depth and the genomic resolution of modifying enzymes, we randomly downsampled both the sequenced molecules and the informative positions within the region of interest (ROI) (Figure 2B). For instance, an informative position density of 10–15% reflects an SMF experiment using M.CviPI, which modifies cytosines in GCH contexts on both strands, whereas a density of 50% corresponds to an experiment using M.EcoGII, which modifies adenines on both strands. Simulations were performed across multiple combinations of experimental parameters, with five to ten random replicates generated for each combination, followed by footprint spectral analysis to estimate the sizes of the footprints present in the data (Figure 2B).

Notably, we observed that the position of peaks in the footprint spectra corresponding to TFs was more sensitive to specific configurations of informative positions than the peak corresponding to non-positioned nucleosomes. This effect was particularly pronounced at lower densities of informative positions and reduced sequencing coverage (Figure 2C).

Intuitively, the spectral analysis for nucleosomes was more robust to random configurations of informative positions because randomly positioned nucleosomes across sequenced molecules spanned a broader range of spacings between informative positions. In contrast, the anchored TF footprint covered only those spacings between informative positions that fell within its specific binding site. Therefore, footprint spectral analysis for short, well-positioned DNA-binding proteins like TFs strongly depends on the density of informative positions and their genomic positions relative to binding sites.

To quantitatively assess the influence of each experimental factor on the performance of the footprint spectral analysis, we conducted a Sobol’ variance-based global sensitivity analysis using the Monte Carlo method (36).

First, we drew 3,072 sets of experimental parameters using Sobol’ sampling from the space of sequencing depth (ranging from 50 to 1,500 molecules), density of informative positions (ranging from 4% to 50%), and background and footprint noise levels (both ranging from 1% to 15%). For each parameter set, we generated five random *in-silico* SMF datasets under the described scenarios (Figures 2A, 2B), resulting in a total of 30,720 *in-silico* datasets. For each dataset, we performed statistical footprint spectral analysis and quantified the model’s performance in recovering ground-truth spectra by calculating the Wasserstein distance (Figure 2D). Finally, we calculated Sobol’ first-order and total effect indices to quantify the contribution of each experimental factor to the variance in the Wasserstein distance (Figure 2E).

Our analysis revealed that the density of informative positions was the most critical factor, accounting for approximately 55%–60% of the total variance in model performance across the two experimental scenarios (Figure 2E). Scatter plots (Figure 2D) showed that the improvement in model performance slowed considerably beyond elbow points of 18.3% and 17.5% for the two scenarios, respectively. Sequencing depth was the second most important factor, explaining 40%–45% of the variance (Figure 2E), with an elbow point around 500–600 sequenced molecules (Figure 2D). In contrast, background and footprint noise levels contributed to only 5%–10% of the variance (Figure 2E).

As our analysis indicated that density of informative positions and sequencing depth contribute the most to the performance of the footprint spectral analysis in recovering ground-truth spectra, we investigated the interaction between these experimental factors (Figure 2F). Notably, this analysis revealed that higher sequencing depth can partially compensate for a low density of informative positions, enabling comparable performance in recovering ground-truth spectra (Figure 2F). For instance, in the scenario with non-positioned nucleosomes, footprint spectral analysis for regions with approximately 6.5% of informative positions achieve similar performance to regions with 16.3% of informative positions when sequencing depth is increased fivefold, from 250 to 1,250 molecules (Figure 2F). Likewise, in the scenario involving both nucleosomes and TFs, the footprint spectral analysis for regions with 8.3% of informative positions achieve similar performance to regions with 21.8% of informative positions when sequencing depth is increased from 250 to 1,250 molecules (Figure 2F).

To contextualize these findings, we analyzed the density of informative positions within 500 bp regions across the mouse (mm10) and human (hg38) genomes for enzymes commonly used in SMF assays, including M.CviPI (GCH), M.EcoGII or Hia5 (adenines), and DddB or SsDddA (cytosines) (Figure 1B). Our analysis revealed that only about 1.2% of regions in the mouse and 2.8% in the human genomes exceeded the 15% cutoff for density of informative positions when using the M.CviPI methyltransferase (Figure 2G). In contrast, close to 100% of 500 bp regions in both genomes had densities exceeding 15% when using M.EcoGII, Hia5, DddB, or SsDddA (Figure 2G).

### A computational model for disentangling footprints with different lengths in SMF datasets

The ultimate goal of any SMF experiment is to detect and classify footprints of DNA-binding protein complexes within individual chromatin fibers. While the footprint spectral analysis described above provides insights into the lengths and abundances of footprints present in the data, it does not specify their exact positions.

To address this, we developed a model that predicts the precise locations of footprints within each molecule for a set of user-defined footprints. Briefly, we determine footprint ranges based on the spectrum obtained from the footprint spectral analysis (Figure 3A) and construct position weight matrices (PWMs) as mathematical representations of footprints in SMF datasets. In these matrices, the alphabet consists of two symbols: 0, representing accessible positions, and 1, representing protected positions (Figure 3B). The number of rows in the PWM corresponds to the footprint length, and each matrix entry represents the probability of observing either a 0 (accessible) or a 1 (protected) at a specific position within a footprint.

**Figure 3:**
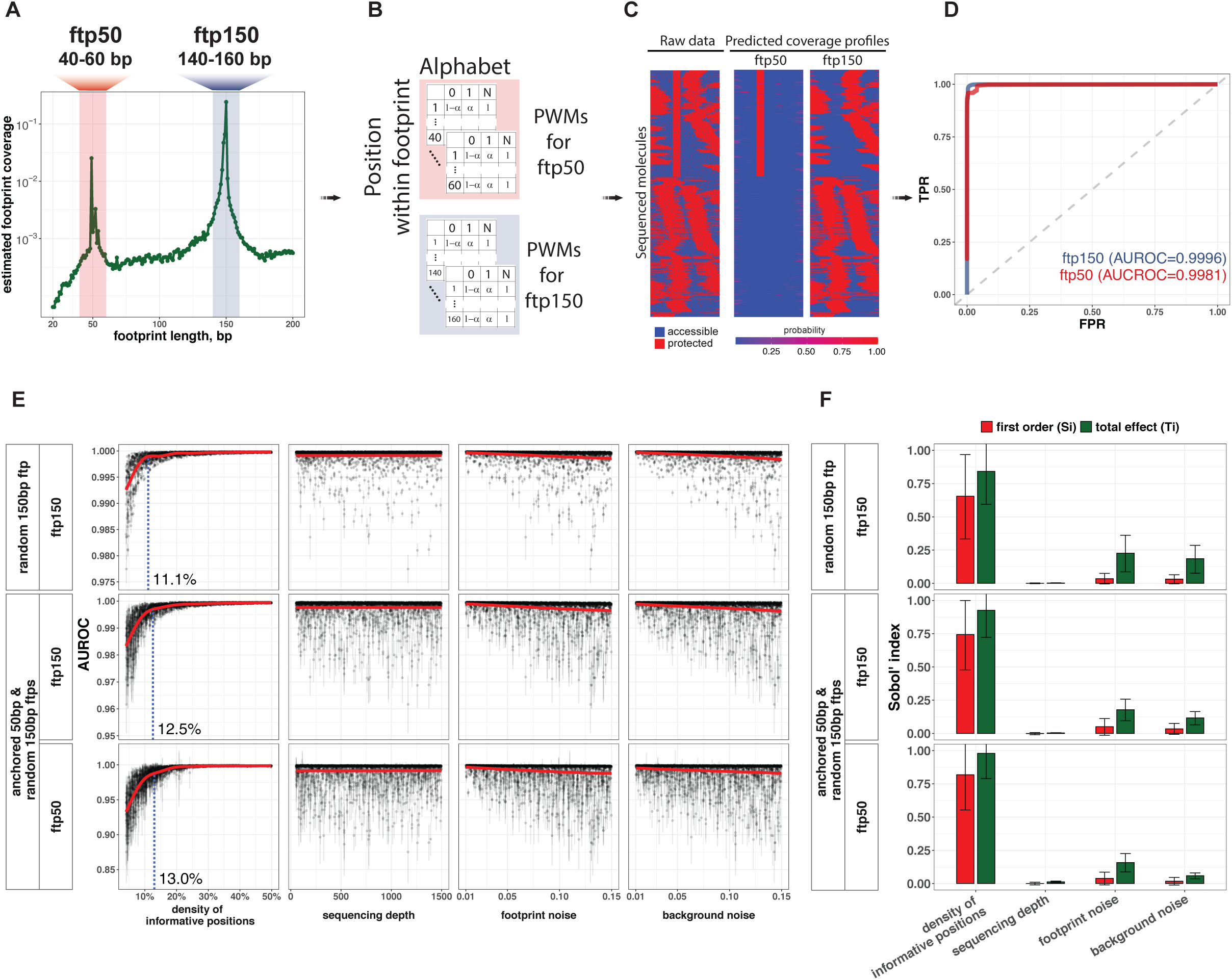
Statistical disentangling of footprints within each molecule in SMF datasets. (A) Example of a footprint spectrum obtained for an *in-silico* generated SMF dataset containing footprints with lengths 50bp (ftp50) and 150bp (ftp150). For prediction of footprint positions we select ranges of footprint lengths around spectral peaks for short (40-60bp for ftp50) and long (140-160bp for ftp150) footprints, sum and renormalize inferred coverages within these ranges and construct position weight matrices in (B). (B) Example of Position Weight Matrices (PWMs) constructed using footprint spectrum from inference results in (A). Each matrix entry represents the probability of observing either a 0 (accessible) or a 1 (protected) at a specific position within a footprint and the number of rows in the PWM corresponds to the footprint length. The third column in the matrices represents non-informative positions (N) and is filled with 1. (C) Example of footprint prediction using the PWMs in (B). Next to the heatmap with raw data, the corresponding predicted coverage for each footprint range is shown. Blue and red colors indicate low (0) and high (1) coverage probabilities respectively. (D) Receiver operating characteristic (ROC) curve for assessing performance of the model at predicting positions covered by short (ftp50 footprint in red) and long (ftp150 footprint in blue) footprints. Areas under the ROC curves (AUROC) are displayed in the inset and used in Sobol’ variance-based global sensitivity analysis for accessing importance of each experimental factor for performance of the model at discriminating footprint positions within SMF datasets. (E) Scatters illustrating performance of the model at discriminating positions covered by different footprints (ftp50 and ftp150) for the two experimental scenarios considered previously (Figure 2A). Y-axes represent areas under the ROC curve as described for (D). Each point and error bars show mean and standard deviation of AUROCs across 5 random *in-silico* SMF datasets. Panels illustrate how AUROCs depend on each experimental factor included in the simulation for two cases described previously in Figure 2A-2B. (F) Sobol’ global sensitivity of the model performance at discriminating positions covered by different footprints to experimental parameters. First order (red) and total (green) Sobol’ indices were calculated for performance of the model at predicting positions covered by ftp150 and ftp50 for each experimental scenario considered in Figure 2A-2B.

Due to limited resolution of enzymes used for marking accessible positions in SMF assays it is very common that, for the vast majority of positions within a sequenced molecule, information regarding their accessibility is missing, resulting in very sparse observed data. To account for missing data, we sum the likelihood across all possible configurations at positions with lacking accessibility information. This approach, in essence, introduces an additional symbol (N) into the alphabet, with all entries in the PWM for this symbol set to 1 (Figure 3B, Supplementary Text Figure S8).

We then use a dynamic programming approach to calculate posterior probabilities for each footprint to cover every position within individual molecules (Figure 3C). This method incorporates carefully defined boundary conditions for partition sums to prevent unwanted edge effects, such as statistical positioning, which can arise when modelling binding of large protein complexes like nucleosomes within relatively short (200-600 bp) sequenced fragments (37) (Supplementary Text, Figure S2, eq. 19).

The prediction of footprints in SMF data enables users to disentangle footprints of varying lengths, such as those corresponding to TFs (ftp50) or nucleosomes (ftp150) (Figure 3C). It also allows for the classification of sequenced chromatin fibers based on their footprint content, such as nucleosome-containing or TF-containing fibers.

Similar to the footprint spectral analysis, we evaluated the model using *in-silico* SMF datasets with known footprint positions and varying experimental factors. We conducted a Sobol’ variance-based global sensitivity analysis to assess the model’s performance in distinguishing between positions covered by long footprints (ftp150), short footprints (ftp50), or background under the two experimental scenarios described previously (Figure 2A).

Briefly, using the footprint spectra obtained from the *in-silico* datasets generated according to Sobol’ sampling of experimental parameters, we constructed PWMs (Figures 3A, 3B) and calculated the probabilities for each position in every molecule being covered by footprints (ftp150 for the first scenario, and ftp50 and ftp150 for the second scenario) (Figure 3C). Using the ground truth from the *in-silico* datasets, we then calculated Areas Under the Receiver Operating Characteristic Curves (AUROC, Figure 3D) to evaluate model performance in detecting positions covered by the respective footprints (Figure 3E).

Finally, we computed first-order and total effect Sobol’ indices to quantify the influence of each experimental factor on the model’s performance in predicting footprint locations (Figure 3F).

Our results reveal that, as with footprint spectral analysis (Figure 2E), the density of informative positions is the most critical experimental factor affecting footprint discrimination in SMF datasets (Figure 3F). Notably, its influence is even more pronounced in this context, with total effect Sobol’ indices ranging from 84–93% of the variance for long footprints (e.g., nucleosomes) to nearly 98% for short footprints (e.g., TFs) (Figure 3F).

In summary, our simulations and Sobol’ sensitivity analysis provide a quantitative evaluation of a key insight: the primary experimental limitation in SMF assays, whether for statistical inference of footprint lengths or for precise localization of different footprints within sequenced molecules, is the enzyme’s ability to provide accessibility information across as many positions as possible. These findings highlight the critical role of enzyme selection in SMF assays, particularly for experiments aimed at detecting TF binding.

### Statistical analysis of SMF experiments carried out with M.CviPI methyltransferase reveals higher methylation rates at nucleosome DNA entry-exit sites

To apply our model to biological SMF data, we designed 48 amplicons targeting regions with distinct chromatin signatures (Supplementary Table 1). Specifically, we included amplicons overlapping binding sites for TFs Ctcf and Rest (4 amplicons for each TF), 15 amplicons representing features typical of active promoters, as well as amplicons targeting gene bodies (1 amplicon), intergenic regions (4 amplicons), bivalent chromatin regions (10 amplicons), and regions characterized by high levels of H3K9me3 and/or DNA methylation, referred to as repressed chromatin (9 amplicons).

We performed Nucleosome Occupancy and Methylome Sequencing (NOMe-seq) experiment on these amplicons in mouse embryonic stem (mES) cells using M.CviPI methyltransferase (20, 38). Nine out of the 48 amplicons yielded fewer than 300 molecules and were excluded from further analysis (Extended Data Figure 1). The remaining 39 amplicons produced an average of 1,450 molecules per amplicon, ranging from 318 to 4,348 molecules. Notably, 49% of the analyzed amplicons (19 out of 39) yielded more than 1,000 molecules each (Figure 4A).

**Figure 4:**
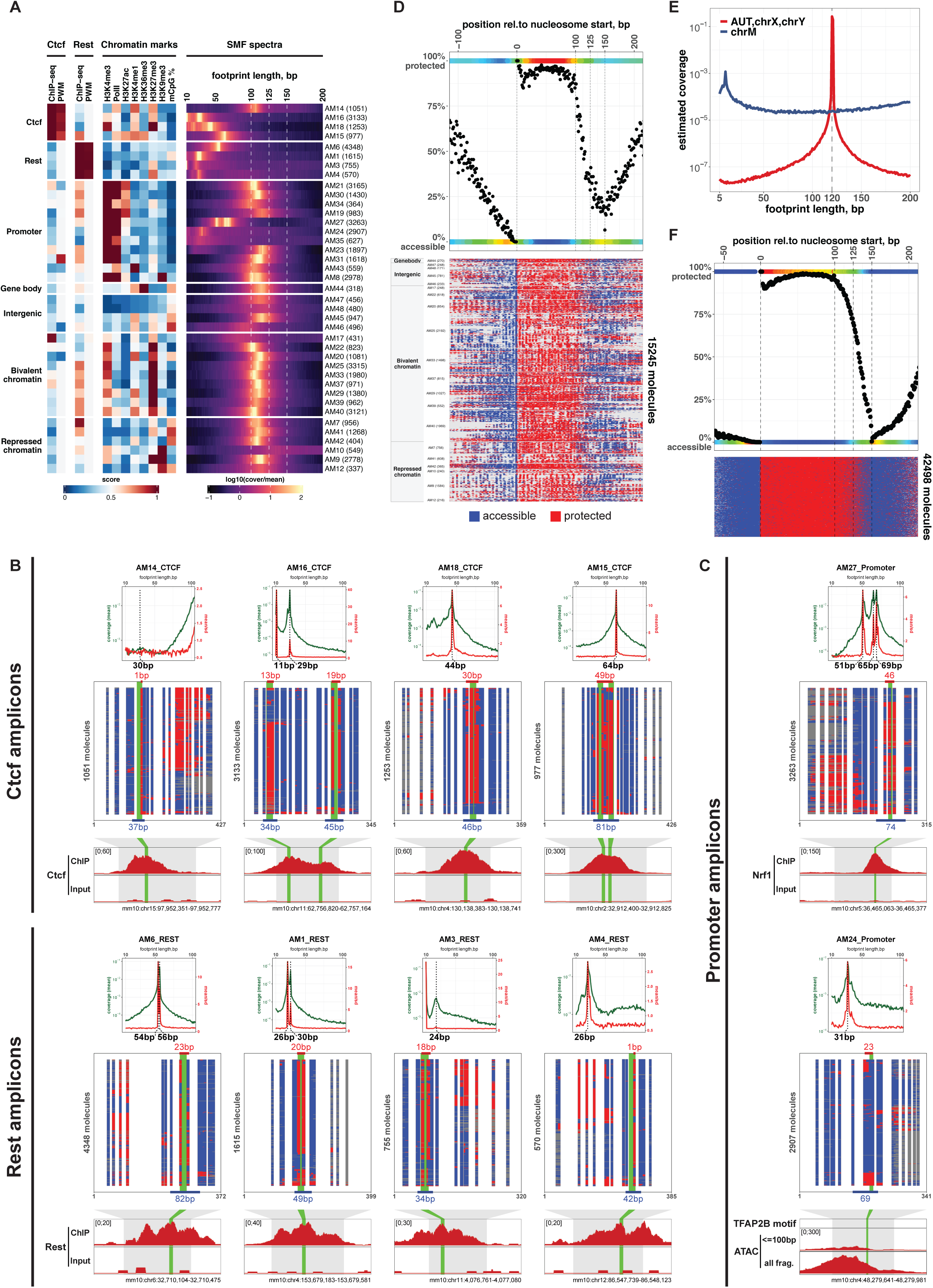
Footprint spectral analysis of NOMe-seq datasets reveals short nucleosome footprints. (A) Heatmaps illustrating chromatin signatures of the designed amplicons as measured by previously published ChIP-seq experiments for TFs Ctcf and Rest (41) as well as for histone post-translation modifications (19) (left panels). The right panels of the heatmaps illustrate results of SMF spectral analyses for corresponding amplicons using datasets from NOMe-seq experiments in mESCs. Amplicon groups based on chromatin signatures, e.g. Ctcf, Rest or Promoter etc., are displayed on the left side of the heatmaps and amplicon names with corresponding number of molecules yielded in the NOMe-seq experiment are shown on the right side of the heatmaps. Only amplicons with number of molecules greater than 300 are included. (B) Footprint spectra (top panels) inferred from NOMe-seq datasets (heatmaps in the middle) for amplicons overlapping Ctcf and Rest binding sites, along with corresponding ChIP-seq and Input profiles (bottom tracks). Spectra are shown for footprint lengths between 10–100 bp, with footprints corresponding to major spectral peaks displayed below each panel. Genomic lengths of TF-like footprints overlapping TF motifs (highlighted in green), as well as regions between flanking accessible GCHs are indicated by red and blue bars at the top and the bottom of each heatmap respectively. (C) Footprint spectra inferred from NOMe-seq datasets for promoter amplicons AM27 and AM24. Annotations for footprint spectra (top panels) and NOMe-seq heatmaps (middle panels) are the same as in (B). Genomic tracks (bottom panels) show ChIP-seq and Input profiles for Nrf1 (AM27), and ATAC-seq profiles for subnucleosomal fragments (<100 bp) and all fragments (AM24). Sequence motifs for NRF1 (AM27) and TFAP2B (AM24) are highlighted in green. (D) SMF data from NOMe-seq experiment aligned around starts of predicted nucleosomes within each molecule (bottom heatmap) and average protection (100 - %GmCH) around nucleosome starts across all molecules (top panel). (E) Footprint spectra for autosomes and sex chromosomes (in red) and mitochondrial DNA (blue) obtained from analysis of a public genome-wide NOMe-seq experiment (29). (F) SMF data obtained from the genome-wide NOMe-seq experiment (29) aligned around starts of predicted nucleosomes (bottom heatmap) and average protection (100 - %GmCH) around nucleosome starts across all molecules (top panel).

We began by performing footprint spectral analysis for each amplicon (Figure 4A, Extended Data Figure 1). As anticipated, the footprint spectra for majority of amplicons targeting TF binding sites (Ctcf and Rest) exhibited distinct peaks corresponding to footprint lengths of 30– 60 bp. However, we observed substantial variability in the estimated footprint lengths across amplicons targeting the same TF (Figure 4A).

To investigate the source of this variability, we annotated motif positions and overlaid NOMe-seq data with ChIP-seq profiles for the corresponding TFs (Figure 4B). In addition, we annotated TF footprints in the NOMe-seq data that overlapped TF motifs (highlighted in red at the top of the heatmaps in Figure 4B) as well as regions bounded by flanking accessible GCH sites (highlighted in blue at the bottom of the heatmaps in Figure 4B).

We found that the footprint lengths estimated by spectral analysis strongly depended on the specific configurations of GCH sites relative to TF motifs. In most cases, the peaks in the footprint spectra fell between the lengths of the protected regions overlapping TF motifs and the distances between flanking accessible GCH sites. These results suggest that the observed variability in footprint length across amplicons targeting the same TF arises from differences in the spatial configuration of informative sites (GCHs) relative to the TF motifs. This observation is consistent with our *in-silico* simulations (Figure 2C) and further underscores the importance of enzyme selection when designing SMF experiments targeting TF binding sites.

Notably, in addition to the amplicons targeting Ctcf and Rest, our footprint spectral analysis identified two promoter-associated amplicons (AM27 and AM24) that exhibited strong peaks for short footprints (Figure 4A). To investigate which TFs might be responsible for these footprints, we searched for TF motifs annotated in the JASPAR database (39) within these regions. Our analysis suggests that the short footprint observed in AM27 likely originates from Nrf1 binding, which is further supported by the Nrf1 ChIP-seq profile generated in mESCs (4), whereas the short footprint in the AM24 amplicon overlaps a sequence motif for TFAP2B (Figure 4C).

Interestingly, many footprint spectra revealed peaks attributable to nucleosomes. While the canonical length of nucleosomal DNA wrapped around the histone octamer is 147 bp (40), our analysis consistently identified nucleosome footprints of 100–120 bp, which are shorter than expected.

To investigate this further, we focused on molecules from amplicon groups expected to contain only nucleosome footprints, i.e. Gene body, Intergenic, Bivalent, and Repressed chromatin regions. We performed footprint prediction using equal prior start probabilities across all footprints with lengths ranging from 100 bp to 150 bp, ensuring no bias toward specific footprint lengths. We aligned nucleosome-containing molecules around the predicted nucleosome start positions (Figure 4D). Analysis of average DNA protection across these nucleosome footprints (top panel in Figure 4D) revealed that regions beyond 120 bp from the start of the nucleosome footprint were accessible to M.CviPI methylation in over 50% of the molecules. Hence, the results confirmed that the majority of nucleosome-like footprints in the NOMe-seq data were indeed shorter than the canonical 147 bp.

To independently validate this observation, we analyzed data from a genome-wide NOMe-seq experiment in mESCs (29). Notably, footprint spectral analysis of molecules aligning to nucleosome-associated chromosomes (autosomes and sex chromosomes) revealed a strong peak at 120 bp, consistent with our amplicon-based findings. In contrast, footprint spectrum for mitochondrial DNA showed no nucleosomal peak, as expected (Figure 4E). Further analysis of nucleosome footprints across 100,000 randomly selected 1-kb genomic regions confirmed that DNA beyond 120 bp from the start of nucleosome footprints exhibited also substantially higher accessibility to M.CviPI methylation (Figure 4F).

In summary, both our analysis of amplicon-based NOMe-seq data (Figure 4D, Extended Data Figure 1) and genome-wide NOMe-seq data (29) (Figure 4E, 4F) identified nucleosome footprints in mESCs to be shorter than the canonical 147 bp suggesting higher accessibility of entry and exit sites to M.CviPI methylation.

### Footprint spectral analysis of SMF dataset at binding sites for TFs in mESCs reveals distinct footprint lengths

We observed that the footprint spectral analysis of amplicons targeting TF binding sites exhibited large variability in estimated footprint lengths across amplicons targeting the same TF due to variability in GCH positioning relative to TF motifs (Figures 4A, 4B). We next investigated whether this limitation can be circumvented by carrying out footprint spectral analysis for large numbers of TF binding sites across the genome.

We analyzed public ChIP-seq datasets for binding of TFs Ctcf and Rest in mESCs and quantified ChIP-seq enrichments at 200bp windows centered around their motifs genome-wide (41). Next, we split motifs for TFs into 4 categories according to their PWM scores and ChIP-seq enrichments, namely 4 combinations of high/low PWM scores and high/low ChIP-seq enrichments (Figure 5A).

**Figure 5:**
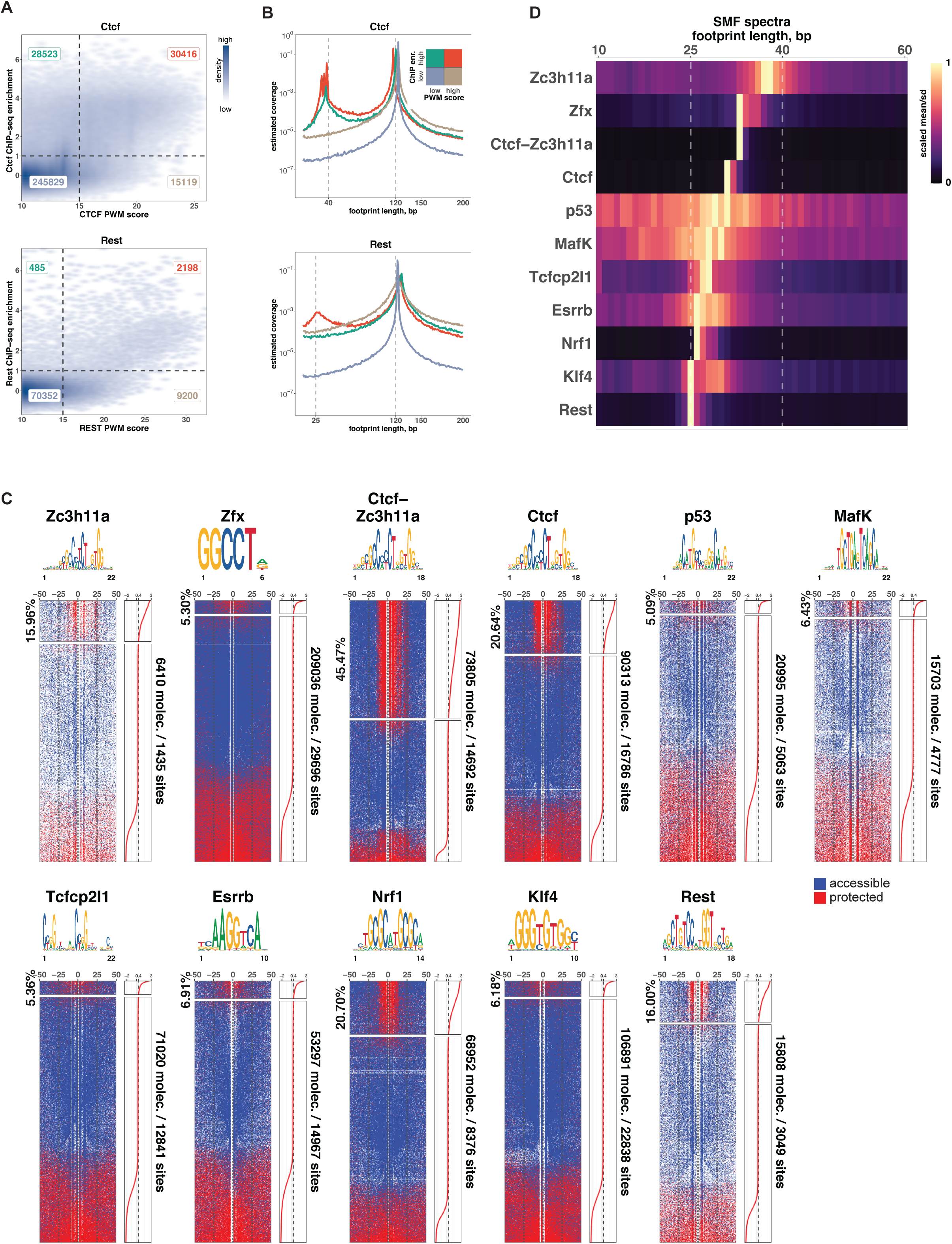
Statistical model reveals variability in footprint lengths for different TFs in mESCs. (A) Two-dimensional densities comparing weight-matrix scores (PWM score) against ChIP-seq enrichments in 200bp windows centered at motifs for TFs Ctcf (top panel) and Rest (bottom panel) (41). Motif containing windows were allocated into 4 groups according to PWM scores and ChIP-seq enrichments for each TF. For each group, number of windows are shown in corresponding quadrants with colors matching colors for resulting footprint spectra in (B). (B) Footprint spectra obtained for the 4 motif groups for TFs Ctcf and Rest defined in (A). (C) Heatmaps illustrating accessibility of cytosines in GCH contexts as measured by NOMe-seq experiment in mESCs (29) at 100bp regions centered around motifs for corresponding TFs. Rows correspond to individual molecules ordered by TF score (shown on the right side of each heatmap) calculated using the model described in this study and averaged across corresponding motif within every molecule. Molecules with TF score above a chosen cutoff (Z-standardized TF score >= 0.4) were classified as containing footprint for TFs and each heatmap was split accordingly, with proportion of TF containing molecules indicated on the left side of each heatmap and total numbers of analyzed molecules and motifs indicated on the right side of each heatmap. (D) Heatmap of footprint spectra derived from NOMe-seq molecules containing high-confidence footprints for TFs (top 10,000 molecules or molecules with TFscore ≥ 0.4) shown in (C). Each row in the heatmap illustrates mean/sd ratios linearly scaled to the interval [0,1] for footprints with lengths from 10 to 60bp.

The subsequent spectral analysis of genome-wide NOMe-seq data (29) revealed footprint spectra with one or two distinct peaks in 400bp regions, that had been centered around the Ctcf or Rest motifs belonging to the 4 described categories (Figure 5B). The larger peak, likely to correspond to a 120 bp nucleosome footprint (Figure 4), was observed in all four categories (Figure 5B). A shorter footprint (∼30-40bp), presumably belonging to the TF Ctcf, was identified for regions with high ChIP-seq enrichments irrespective of the Ctcf PWM score. For regions containing the Rest motif, however, a short footprint (∼25bp) was only identified for regions with high ChIP-seq enrichments and having a strong Rest PWM score, possibly due to lower number of regions with high ChIP-seq enrichments and low Rest PWM score (Figure 5A, 5B).

To assess the variability of footprint lengths among different TFs we extended our analysis to a broader panel of TFs. We used annotations from a previous study (42) that analyzed ChIP-seq enrichments obtained from 14 public ChIP-seq datasets (4, 43–49) at 201bp regions centered around corresponding TF motifs. We selected regions with high ChIP-seq enrichments (log2(ChIP/Input) ≥ 1) and having at least one GCH within a 30bp window centered around the TF motif (Extended Data Figure 2A) and retrieved all molecules from genome-wide NOMe-seq datasets for mESCs (29) overlapping the selected motifs.

We noticed that some groups of TFs have substantial overlaps of their binding regions (Extended Data Figure 2B). Therefore, we split regions for TFs that have significant overlaps with each other, e.g. Nanog, Sox2 and Oct4, into groups of regions where all TFs co-occur, e.g. Nanog-Sox2-Oct4, and regions where binding of only one of the TF has been observed, e.g. Nanog, Sox2 and Oct4 (Extended Data Figure 2B). Similarly, we split regions of Ctcf and Zc3h11a into groups with both TFs having high ChIP-seq enrichments and regions where only Ctcf or Zc3h11a binding has been observed.

We used our statistical model to predict shorter (15-80bp, TF footprint) and longer (100-150bp, Nucleosome footprint) footprints in each molecule overlapping with resulting groups of regions and ordered all molecules according to predicted TF score (Figures 5C, Extended Data Figure 2C, see Methods). Finally, we selected molecules with high-confidence footprints for TFs (top 10,000 molecules or molecules with TFscore >= 0.4 (Figures 5C, Extended Data Figure 2C) and performed footprint spectral analysis to estimate footprint lengths for each TF (Figures 5D, Extended Data Figure 2D). Our footprint spectral analysis revealed distinct footprint lengths for 11 out of 16 groups for TF binding which vary between ∼25bp and ∼40bp (Figures 5D, Extended Data Figure 2D). Taken together, our analysis of public ChIP-seq datasets in mESCs integrated with analysis of public genome-wide SMF datasets using our statistical model reveals distinct footprint lengths at sequence motifs for TFs in mESCs.

### *De novo* identification of TF binding sites in genome-wide SMF datasets

The efficient packaging of eukaryotic DNA is facilitated by its wrapping around histone octamers. However, this structural organization presents challenges for the *de novo* identification of TF binding sites in single-molecule footprinting (SMF) assays, as most observed footprints result from DNA protected by nucleosomes (Figure 4D-4F). While previous studies have proposed methods to identify TF footprints in SMF datasets, these approaches often rely on prior knowledge of TF binding sites, such as known motifs or ChIP-seq enrichment data (50–52). Our statistical model addresses this limitation by enabling the rigorous calculation of posterior probabilities for footprints of distinct lengths, effectively disentangling TF footprints from nucleosomal ones (Figure 3B-3C).

To evaluate the model’s performance in detecting regions with consistently high occurrences of accessible positions and TF footprints across multiple molecules, we selected 11 genomic regions ranging from 1 to 1.5 Mb in length, covering a total of approximately 12.8 Mb. These regions were chosen based on their high densities of ATAC-seq peaks (data from (53)) and/or the presence of binding sites for Ctcf and Rest (data from (41)) in mESCs. From the genome-wide NOMe-seq dataset in mESCs (data from (29)), we selected all molecules with densities of GCHs exceeding 5% and longer than 50 bp overlapping the 11 genomic regions. For each molecule, we calculated posterior probabilities for shorter TF footprints (15–80 bp) and longer nucleosome footprints (100–150 bp) (see Methods). Based on these probabilities, we derived scores for free accessible positions and TF footprints and chose cutoffs to identify high-confidence accessible positions (background, BG) and those likely covered by TFs (see Methods).

Using sliding windows of 500 bp with a 250 bp step size, we quantified the number of accessible (BG) and TF positions across all overlapping molecules. We then applied a binomial test to assess the statistical significance (adjusted for multiple comparisons using Benjamini & Hochberg (FDR) method) of the observed fractions of accessible (Extended Data Figure 3A) and TF (Extended Data Figure 3B) positions, testing the alternative hypothesis that these fractions are higher than expected fractions across all molecules overlapping selected regions (Figure 6A). Interestingly, 39% (1,891 out of 4,789, Extended Data Figure 3A) of sliding windows with significantly higher frequencies (BG FDR ≤ 1%, −log10(FDR) tracks in Figure 6A, Extended Data Figure 3D) of accessible positions overlapped with high-accessibility regions identified by ATAC-seq experiments in mESCs (data from (53)).

**Figure 6:**
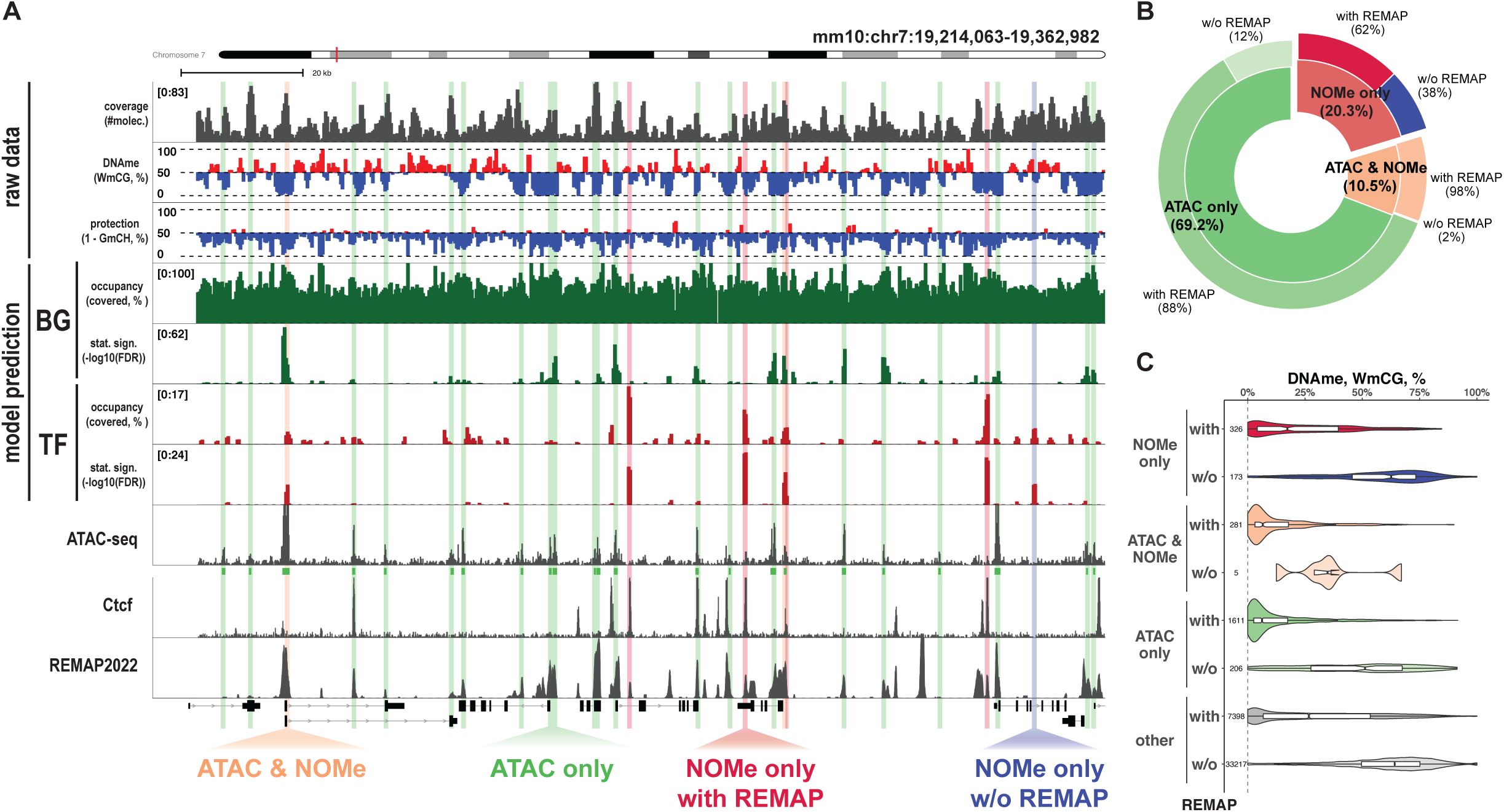
Masking nucleosomal footprints allows better interpretation of SMF datasets. (A) Genome browser tracks illustrating raw NOMe-seq data for mESC (29), predicted occupancy and statistical significance for background (BG, accessible regions) and TF footprints (TF, predicted footprints with lengths from 15 to 80bp), ATAC-seq enrichments and peaks (53), Ctcf ChIP-seq enrichments (41) and coverage of ChIP-seq peaks in mESCs annotated in REMAP2022 database (54). All tracks illustrate values for sliding windows of 500bp with a 250bp step. Tracks with raw SMF data show coverage of each window by NOMe-seq reads (track coverage), proportion of methylated WCGs (track DNAme) and protected (unmethylated) GCHs (track protection) across all molecules overlapping each window. Tracks with model prediction illustrate estimated proportions of GCHs categorized as accessible (BG tracks) and covered by TF footprints (TF tracks) across all molecules overlapping each window and corresponding statistical significances (-log10(FDR) as calculated using binomial test and adjusted for multiple testing). Highlighted regions indicate windows with significant enrichment of TF footprints (FDR ≤ 0.01) and overlapping ATAC-seq summit (“ATAC & NOMe”, in orange), containing only ATAC-seq summit (“ATAC only”, in green) or having only significant enrichment of TF footprints confirmed by having ChIP-seq summit in REMAP2022 (“NOMe only with REMAP”, in red) or lacking REMAP2022 ChIP-seq summit (“NOMe only w/o REMAP”, in blue). (B) Pie chart summarizing percentages of sliding windows belonging to each group (“ATAC only”, “ATAC & NOMe”, “NOMe only”) as well as fraction of windows within each group that overlap ChIP-seq summit annotated by REMAP2022 database in mESCs. (C) Levels of WCG methylation at sliding windows across all 11 genomic loci selected in this study belonging to each group and split by overlap with ChIP-seq summit annotated by REMAP2022 database in mESCs. “other” indicate windows which were not classified as “ATAC only”, “ATAC & NOMe” or “NOMe only”.

Next, we assessed the performance of our model to detect genomic windows enriched for TF footprints. We identified 852 such windows (FDR ≤ 1%) out of 48,862 within the 11 selected regions (∼1.7%, Extended Data Figure 3B, 3C) and examined their overlap with ATAC-seq summits and ChIP-seq summits annotated in the REMAP2022 database (54) (Figure 6B). These windows were classified into three groups: (i) those overlapping both ATAC-seq summits and having significant TF footprint enrichments in NOMe-seq (“ATAC & NOMe”), (ii) those overlapping only ATAC-seq summits (“ATAC only”), and (iii) those with significant TF footprint enrichments in NOMe-seq but having no overlapping ATAC-seq peaks (“NOMe only”) (Figure 6B, Extended Data Figure 3C). The majority of windows (69.2%, 1,913 windows) fell into the “ATAC only” category, while 10.5% (290 windows) were supported by both assays (“ATAC & NOMe”), suggesting that the relatively low resolution of NOMe-seq limits its ability to detect TF footprints in regions with sparse GCH coverage. Nevertheless, our model identified 562 “NOMe only” windows (20.3%), and 62% of these (351 windows) overlapped previously annotated ChIP-seq summits (Figure 6A–6B, Extended Data Figure 3C), supporting the biological relevance of these predictions.

We further examined endogenous DNA methylation by leveraging CpG methylation information from the NOMe-seq dataset. We observed that regions significantly enriched for accessible positions (as predicted by our model) exhibited strong hypomethylation (Extended Data Figure 3A). To further assess the reliability of TF footprint predictions, we compared DNA methylation levels across the three window groups described above, split by overlap with REMAP2022 ChIP-seq summits (Figure 6C). Consistent with previous observations (48), we found that TF binding frequently occurs in low-methylated regions (LMRs). Regions from the “NOMe only”, “ATAC & NOMe”, and “ATAC only” groups that overlapped ChIP-seq summits exhibited similarly low CpG methylation levels. In contrast, windows without ChIP-seq support showed higher methylation levels, but these levels were comparable between “NOMe only” and “ATAC only” groups (Figure 6C), suggesting that some DNA-binding complexes may occupy regions with higher DNA methylation.

In conclusion, our statistical model effectively disentangles footprints of different lengths, allowing for the masking of nucleosome footprints in SMF datasets, which facilitates the *de novo* identification of regions with accessible chromatin and TF binding sites, including sites that were missed by ATAC-seq assay but are confirmed by direct ChIP-seq experiments (Figure 6A-6B, Extended Data Figure 3C-3D).

## DISCUSSION

Single molecule footprinting assays offer unique opportunities for studying genome regulation by providing high-resolution insights into protein-DNA interactions, chromatin accessibility, and the dynamic organization of nucleosomes and TFs at the level of individual DNA molecules. In this study, we introduced a novel statistical framework for the quantitative analysis of SMF datasets, addressing two main questions that arise during such analyses.

Firstly, our statistical approach allows the unbiased inference of footprint lengths represented in the dataset, accounting for experimental noise. Using a count table that represents the number of observed combinations of accessible and protected positions at various distances, our model generates a footprint spectrum. This spectrum provides information about the abundances of footprints within a certain range and the model can optionally estimate noise levels within accessible and protected positions.

Secondly, employing a dynamic programming approach with carefully defined boundary conditions, our model rigorously calculates the probabilities for each footprint at every position within each SMF molecule, considering noise levels for accessible and protected positions. This process effectively disentangles footprints of different lengths and facilitates the biological interpretation of SMF datasets.

We tested our statistical framework using both *in-silico* and biological SMF datasets. Our variance-based sensitivity analysis underscores the importance of experimental factors such as the density of informative positions, determined by the specific enzyme used in the SMF experiment, and the sequencing depth for the feasibility of statistical analysis.

Our analysis of datasets generated from mESCs by the NOMe-seq assay using M.CviPI yielded interesting biological insights. Specifically, we observed that nucleosome footprints estimated by NOMe-seq assays are, on average, shorter than the canonical ∼147 bp footprint observed in classical biochemical MNase digestion studies (55, 56). This likely reflects higher accessibility of nucleosomal DNA at entry-exit sites due to nucleosome breathing dynamics (55, 57–61). Additionally, our analysis of footprints at TF binding sites revealed various distinct footprint lengths, highlighting the model’s sensitivity in detecting diverse protein-DNA interactions.

A significant feature of our approach is its ability to disentangle footprints of different lengths while accounting for experimental noise in SMF datasets. This capability allows for masking prevalent nucleosome footprints, facilitating the identification and analysis of regions with open chromatin and TF binding without prior information about their locations. Our analysis of NOMe-seq data generated in mESCs demonstrates that, despite the low genomic resolution and sequence bias of M.CviPI, our method reliably identifies regions enriched for TF footprints, many of which are supported by ChIP-seq data but are undetectable using ATAC-seq assay.

Our model relies on detecting enzymatic modifications on DNA, from GCH methylation states from NOMe-seq assays using M.CviPI, adenine methylation states from Fiber-seq assays using Hia5/M.EcoGII, to cytosine deamination states from FOODIE/DAF-seq assays using DddB/SsDddA enzymes. This universality makes our statistical approach adaptable to any SMF assay that provides binary accessibility information at the single-molecule level. The accuracy of the framework depends on the resolution and precision of the specific enzyme and sequencing platform used to mark and detect accessible versus protected positions. Consequently, we anticipate improved performance in identifying regions of open chromatin and TF binding when applying our model to datasets generated with higher-resolution assays such as Fiber-seq (25, 26, 34), FOODIE (27) or DAF-seq (28).

In summary, our statistical framework enables the quantitative analysis and interpretation of SMF datasets. To do so, we developed the R package *nomeR* for analyzing SMF datasets using the statistical approach described in this study, as well as the R package *fetchNOMe* for the fast retrieval of data structures required for the statistical analysis from BAM files generated in NOMe-seq experiments.

---

**Extended Data Figure 1:**
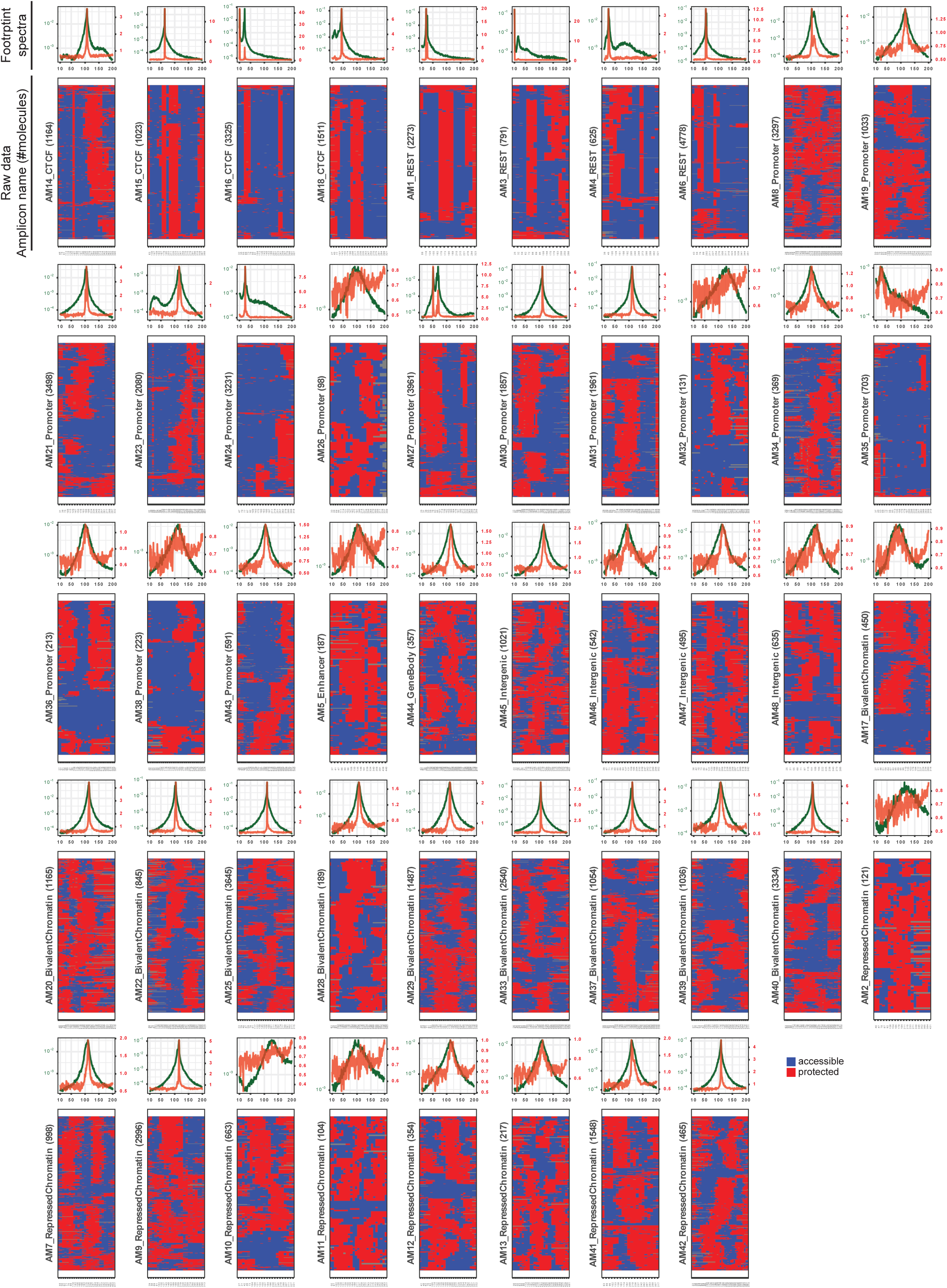
Raw data and footprint spectral analysis from amplicon-based NOMe-seq experiment in mESCs. Raw NOMe-seq data for 48 designed amplicons and corresponding footprint spectra. Each heatmap shows reduced representation of SMF data where rows correspond to individual molecules and only informative positions are shown. Colors of the heatmap depict protection (red) or accessibility (blue) at informative positions. Green and red curves on top of every heatmap illustrate two alternative representations of corresponding footprint spectra. Green curve represents mean values and red curve represents ratios between means and standard deviations across samples from mean-field variational approximation of posterior distribution for footprint coverages.

**Extended Data Figure 2:**
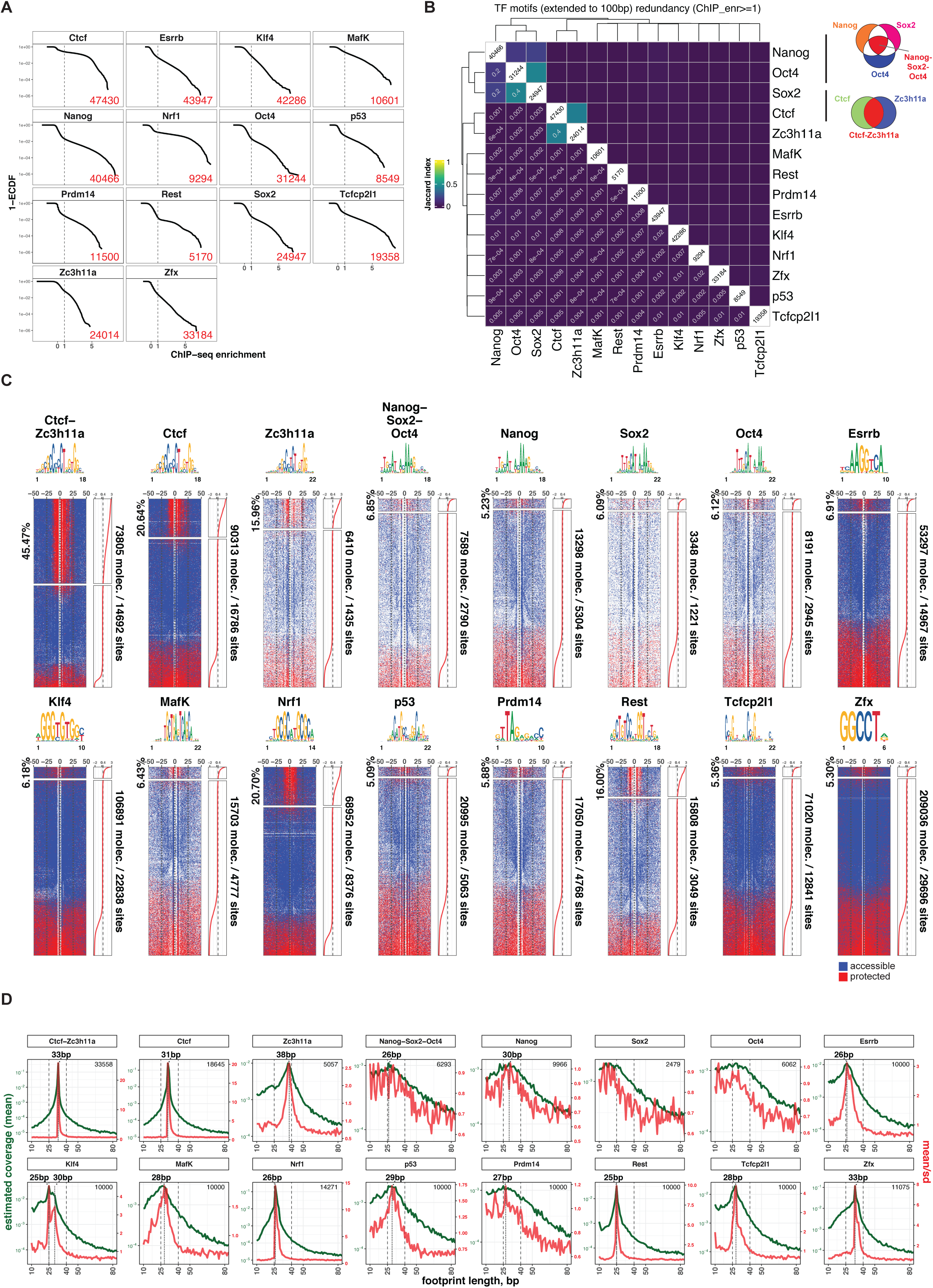
Statistical analysis of NOMe-seq molecules at TF binding sites. (A) Reverse empirical cumulative distributions of ChIP-seq enrichments for TFs at 200bp regions centered around corresponding sequence motifs. Numbers of regions remained after applying a cutoff for ChIP-seq enrichments (log2(ChIP/Input) >= 1) are shown in red in each panel. (B) Heatmap illustrating pairwise co-occurrence of TF motifs with enriched ChIP-seq signal. Color represents Jaccard index for overlaps of 100bp regions centered around selected TF motifs. Motifs for TFs that show substantial overlap were divided into subgroups based on occurrences of all motifs or only individual motifs. In particular, motifs for Nanog, Sox2 and Oct4 were divided into 4 sub-groups: with all three TFs in proximity to each other, or only individual TF within 100bp region having ChIP-seq enrichment above chosen cutoff (log2(ChIP/Input >= 1). Similarly, motifs for Ctcf and Zc3h11a were split into three groups based on co-occurrence of the TFs. (C) Heatmaps illustrating accessibility of cytosines in GCH contexts as measured by NOMe-seq experiment in mESCs (29) at 100bp regions defined in (B) and centered around motifs for corresponding TFs. Rows in the heatmaps correspond to individual molecules ordered by TF score calculated using the model described in this study and averaged across corresponding motif within every molecule. Molecules with TF score above a chosen cutoff (Z-standardized TF score >= 0.4) were classified as containing footprint for TFs and each heatmap was split accordingly, with proportion of TF containing molecules indicated on the left side of each heatmap and total numbers of analyzed molecules and motifs indicated on the right side of each heatmap. (D) Footprint spectra obtained after statistical spectral analysis of selected molecules in (C) at 100bp regions centered around motifs for corresponding TFs (see Methods). Green and red curves illustrate two alternative representations of corresponding footprint spectra. Green curve represents mean values and red curve represents ratios between means and standard deviations across samples from mean-field variational approximation of posterior distribution for footprint coverages. Number of molecules used for statistical footprint spectral analysis is shown in every panel.

**Extended Data Figure 3:**
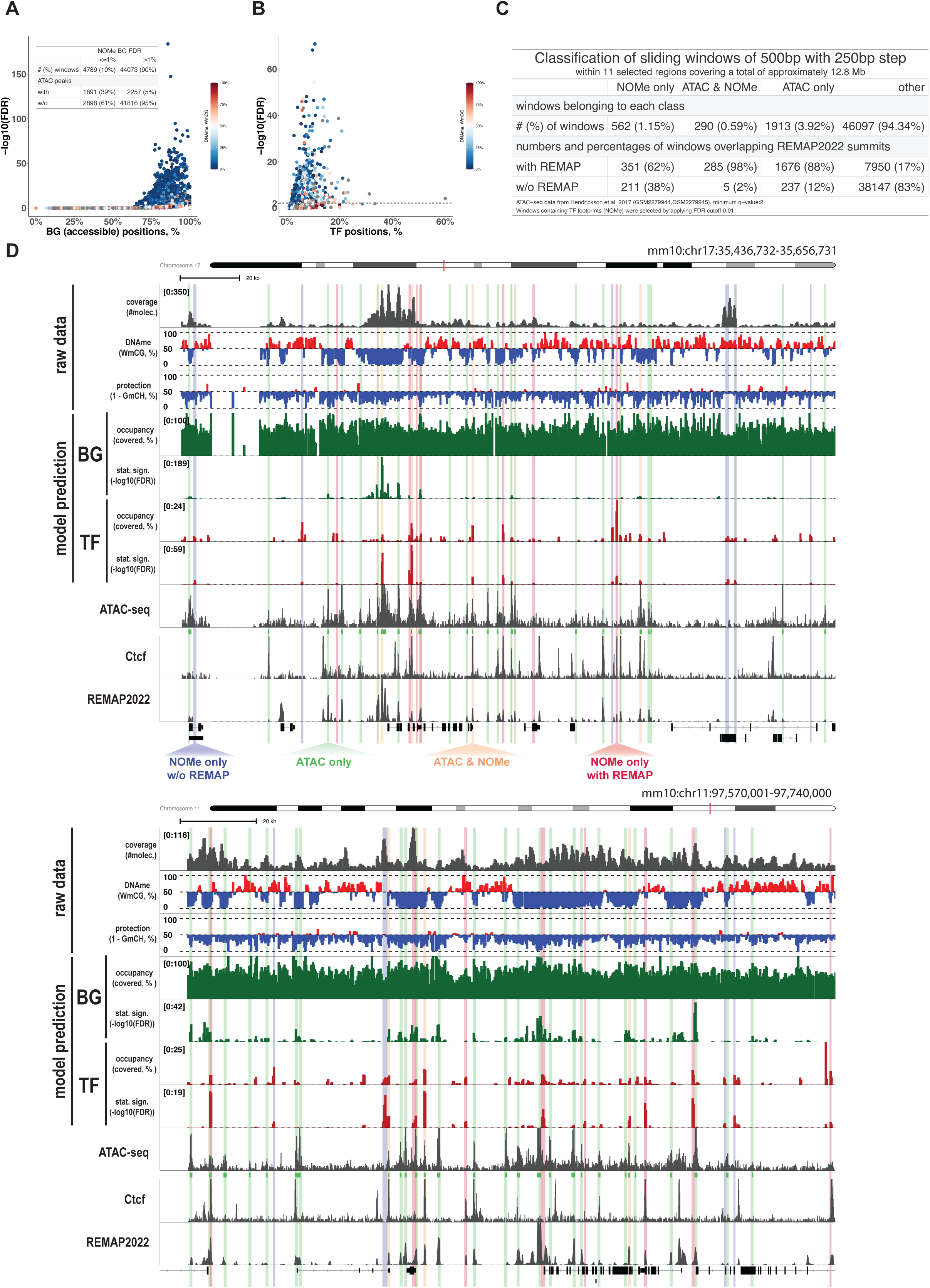
*De novo* identification of regions significantly enriched for accessible sites and TF-protected positions. (A) Volcano plot for genomic sliding windows of 500bp with a 250bp step for detecting regions enriched for occurrences of accessible positions. X-axis represents estimated percentage of accessible positions across all molecules overlapping genomic sliding windows and Y-axis represents −log10 of False Discovery Rates as calculated using binomial test with alternative hypothesis that observed fraction of accessible positions is higher than average fraction across all analyzed molecules and adjusted for multiple testing using Benjamini-Hochberg method. Every point colored by average WCG methylation across all molecules overlapping corresponding genomic sliding window. An inset table shows number and percentages of sliding windows with (FDR ≤ 1%) and without (FDR > 1%) statistically significant enrichment of accessible positions and overlapping (with) or not overlapping (w/o) ATAC-seq peaks. (B) Volcano plot, similar to (A), but for detecting regions enriched for occurrences of TF footprints. X-axis represents an estimated percentage of positions covered by TF footprints across all molecules overlapping genomic sliding windows and Y-axis represents −log10 of False Discovery Rates. Every point colored by average WCG methylation across all molecules overlapping corresponding genomic sliding window. (C) Table illustrating numbers and percentages of genomic sliding windows categorized by statistically significant enrichment (FDR ≤ 1%) of TF footprints as identified by our model and overlaps with ATAC-seq summits and ChIP-seq summits as annotated in REMAP2022 database for mESCs. “ATAC & NOMe” and “NOMe only” represent windows with statistically significant enrichment of TF footprints with and without overlaps with ATAC-seq summits respectively, whereas “ATAC only” represent windows that have overlaps with ATAC-seq summits but lack enrichments of TF footprints according to NOMe-seq data. Class “other” represents all other windows within 11 selected regions which did not fall into the three described groups. Rows “with REMAP” and “w/o REMAP” represent numbers and percentages of windows in each group with and without overlapping ChIP-seq summits annotated in REMAP2022 database for mESC (54). (D) Genome browser tracks for two genomic regions illustrating raw NOMe-seq data for mESC (29), predicted occupancy and statistical significance for background (BG, accessible regions) and TF footprints (TF, predicted footprints with lengths from 15 to 80bp), ATAC-seq enrichments and peaks (53), Ctcf ChIP-seq enrichments (41) and coverage of ChIP-seq peaks in mESC annotated in REMAP2022 database (54). All tracks illustrate values for sliding windows of 500bp with a 250bp step. Tracks with raw SMF data show coverage of each window by NOMe-seq reads (track coverage), proportion of methylated WCGs (track DNAme) and protected (unmethylated) GCHs (track protection) across all molecules overlapping each window. Tracks with model prediction illustrate estimated proportions of GCHs categorized as accessible (BG tracks) and covered by TF footprints (TF tracks) across all molecules overlapping each window and corresponding statistical significances (-log10(FDR) as calculated using binomial test and adjusted for multiple testing using Benjamini-Hochberg method). Highlighted regions indicate windows with significant enrichment of TF footprints (FDR ≤ 0.01) and overlapping ATAC-seq summit (“ATAC & NOMe”, in orange), containing only ATAC-seq summit (“ATAC only”, in green) or having only significant enrichment of TF footprints confirmed by having ChIP-seq summit in REMAP2022 (“NOMe only with REMAP”, in red) or lacking REMAP2022 ChIP-seq summit (“NOMe only w/o REMAP”, in blue).

**Supplementary Table 1**

Annotation for amplicons designed for the NOMe-seq experiment for mESC carried out in this study.

## MATERIALS AND METHODS

### Statistical footprint spectral analysis

Footprint spectral analysis is based on the statistical inference of parameters for a stochastic process, described in detail in the Supplementary Text. Briefly, single-molecule footprinting (SMF) datasets are modelled using a stochastic process that randomly places 𝐾 footprints of lengths 𝒍 = (𝑙_1_, …, 𝑙_𝐾_) on an infinite sequence, with prior start probabilities 𝝅 = (𝜋_1_, …, 𝜋_𝐾_) reflecting their abundances. The index 1 is reserved for accessible positions (i.e., background), where 𝑙_1_ = 1 and 𝜋_1_ represents the prior probability of placing an accessible position during the stochastic process. The lengths (𝑙_2_, …, 𝑙_𝐾_) are user-defined.

We assume that footprints, including the background, cannot overlap and do not interact with each other. Both background and footprints emit either 0 (unprotected position) or 1 (protected position) with specific emission probabilities that reflect noise levels. The emission probability of a protected position within the background is denoted as 𝛽^(1)^, while 𝛼^(1)^ represents the emission probability of a protected position within footprints (see Figure S1 in the Supplementary Text).

For the described stochastic process, we derived the joint probability 𝑃(𝛾, 𝜃|𝑆) of observing a pair 𝛾, 𝜃 ∈ {0,1} representing combinations of accessible (encoded as 0) and protected (encoded as 1) positions at a distance 𝑆 base pairs apart (see Equation 36 in the Supplementary Text). Using these joint probabilities, we constructed a Bayesian model (see Figure S6B in the Supplementary Text) to infer the parameters 𝝆 = (𝜌_1_, 𝜌_2_, …, 𝜌_𝐾_), where 𝜌_1_ represents the coverage of the background, and (𝜌_2_, …, 𝜌_𝐾_) represent coverages for the respective footprints. Additionally, the model infers 𝛽^(1)^ and 𝛼^(1)^, which reflect the probabilities of emitting protected positions within the background and footprints, respectively. These parameters are estimated from the observed frequencies 𝑁^(𝛾θ)^ at various distances 𝑆𝑆 in the SMF dataset (Figure 1D).

Throughout the manuscript, we refer to the vector 𝝆_−𝟏𝟏_ = (𝜌_2_, …, 𝜌_𝐾_) as the footprint spectrum, 𝛽^(1)^ as the background noise, and 1 − 𝛼^(1)^ as the footprint noise.

The Bayesian model was implemented in the R package *nomeR* using the *Stan* programming language (62), and parameter inference was performed using mean-field variational approximation (63) via the *infer_footprints_vb* function. The *nomeR* package supports three possible modes of statistical inference, controlled by parameter *ftp_bg_model.* Setting *ftp_bg_model* to “informative_prior” enables the model to infer both 𝛽^(1)^ and 𝛼^(1)^ in addition to the footprint spectrum. When set to “bg_fixed”, the model assumes that 𝛽^(1)^ is known and constant, while “ftp_bg_fixed” assumes that both 𝛽^(1)^ and 𝛼^(1)^ are known and fixed.

In this study, we performed footprint spectral analysis with ftp_bg_model set to “informative_prior”.

### Predicting footprint positions in single molecule footprinting datasets

Given reasonable estimates of footprints lengths, their abundances (either 𝝆 or 𝝅) and emission probabilities (𝛼^(1)^ and 𝛽^(1)^), our model can predict the locations of footprints within each single molecule in SMF datasets.

Briefly, we construct positional weight matrices (PWM) as mathematical representations for each footprint expected in the SMF dataset. In these matrices (Figure 3B), columns represent an alphabet consisting of 0 (accessible positions), 1 (protected positions) and N (non-informative positions), while rows correspond to positions within the footprint. Each matrix entry represents the probability of observing a 0 or 1 at a specific position within a footprint. The column corresponding to non-informative positions (N) is filled with a probability of 1, as a result of marginalizing the likelihood over all possible variants for unobserved data (see Supplementary Text for details).

Using a dynamic programming approach with the constructed PWMs, we consider all possible combinations of footprint placements and calculate the posterior probabilities for each footprint to start at, or cover, every position within each molecule. Importantly, we reformulated the model and derived asymptotic boundary conditions for partition sums, assuming an infinite sequence without positional preferences for footprints. This adjustment allows us to avoid unwanted boundary effects, such as statistical positioning (37) which can distort posterior probability predictions, especially in short sequenced fragments (500–600 bp) (see Supplementary Text for details).

### *In-silico* simulation of SMF datasets and sensitivity analysis

#### Generation of in-silico datasets

Configurations of non-overlapping footprints and accessible positions along a 10 kb chromosome were randomly generated for 3,000 chromatin fibers, with a 500 bp region of interest (ROI) located at the center of the chromosome (positions 5001–5500). Two experimental scenarios, as described in the main text (Figure 2A), were considered. Both scenarios include a long (150 bp) footprint representing nucleosomes, which lacks positional preferences and covers 50% of the positions within the ROI. In the second scenario, an additional short (50 bp) footprint representing TFs was introduced. This footprint competes with nucleosome footprints, covers 5% of the positions, and occupies a specific site within the ROI. The generation of *in-silico* SMF datasets was implemented in the R package *nomeR* using the generate_insilico_SMF_data function.

To simulate SMF datasets with specified experimental parameters, the following steps were performed:

1. Random mutation of data points within accessible (background) positions according to the specified background noise level.
2. Random mutation of data points within footprints according to the specified footprint noise level.
3. Random sampling of chromatin fibers based on the desired sequencing depth.
4. Random selection of informative positions within the ROI based on the specified density of informative positions.

These procedures were applied to 3,000 *in-silico* chromatin fibers for two experimental scenarios across various combinations of experimental parameters to generate the SMF datasets. For each set of experimental parameters we generated 10 (Figure 2C) or 5 (Figures 2D-2F) random *in-silico* SMF datasets.

For the Sobol’ global variance-based sensitivity analysis using Monte Carlo method we generated 3,072 sets of experimental parameters using function *sobol_matrices* from the R package *sensobol* (36). The experimental parameters varied within the following ranges:

1. Sequencing depth: 50 to 1,500 molecules
2. Density of informative positions: 4% to 50% of positions within the ROI
3. Background noise (𝛽^(1)^): 1% to 15% of accessible positions
4. Footprint noise (1 − 𝛼^(1)^): 1% to 15% of positions covered by footprints.

For each set of experimental parameters, we generated 5 random SMF datasets for the two experimental scenarios described above, resulting in a total of 30,720 *in-silico* datasets. Since our simulations are stochastic but Sobol’ sensitivity analysis is designed for deterministic functions, we addressed this discrepancy by setting a fixed seed for the pseudo-random number generator in each round of SMF data generation, ensuring reproducibility of the random process.

#### Sobol’ sensitivity of the footprint spectral analysis

To assess the influence of experimental factors on footprint spectral analysis, we performed footprint spectrum inference using the *infer_footprints_vb* function from the *nomeR* package for each *in-silico* SMF dataset generated for the Sobol’ sensitivity analysis. The inferred footprint spectra were then compared to the ground-truth spectra corresponding to each experimental scenario by calculating the Wasserstein distance using the *wasserstein1d* function from the R package *transport* (64).

To evaluate the sensitivity of model performance to experimental parameters, as measured by the Wasserstein distance to the ground-truth spectra, we first averaged the Wasserstein distances across the five random datasets for each combination of experimental parameters. Scatters in Figure 2D illustrate mean and standard deviation of the Wasserstein distances across random datasets for the same set of experimental parameterss. Smoothed curves were obtained by fitting generalized additive model using the *gam* function from the R package *mgcv* (65) and elbow points were estimated using the R package *kneedle* (66).

The averaged values of the Wasserstein distances were then used as input for the *sobol_indices* function from the R package *sensobol* (36) to calculate both first-order and total Sobol’ indices (Figure 2E).

To investigate interaction between the density of informative positions and the sequencing depth in determining the performance of the footprint spectral analysis (Figure 2F) we used a generalized additive model (*gam* function from the *mgcv* R package) to approximate Wasserstein distances on a regular 200 x 200 grid for the two experimental parameters and illustrated this approximation using a heatmap with contour lines obtained by the function *geom_contour* from the ggplot2 R package (67) with default parameters.

#### Sobol’ sensitivity of footprint prediction

Similarly, we assessed the influence of experimental factors on the precision of footprint position predictions within *in-silico* SMF datasets. Specifically, based on the inferred footprint spectra and noise levels, we calculated coverage probabilities for short (40–50 bp, “ftp50”) and long (140–160 bp, “ftp150”) footprints for all informative positions within each molecule.

To evaluate the performance of these predictions, we used predicted footprint coverage probabilities and the ground-truth positions of footprints to calculate the area under the receiver operating characteristic curve (AUROC, Figure 3D) using the R package *pROC* (68).

Finally, the AUROC values, averaged across the five random datasets for each combination of experimental parameters, were used as input for the *sobol_indices* function from the R package *sensobol* (36) to calculate both first-order and total Sobol’ indices (Figure 3F). Scatters, smoothed curves and elbow points for AUC-ROC (Figure 3E) were obtained similarly as described for the sensitivity of the footprint spectral analysis.

#### Densities of informative positions for various SMF assays in mouse and human genomes

To estimate the densities of informative positions for SMF assays within 500 bp regions of the mouse (*mm10*) and human (*hg38*) genomes, we selected 500 bp tiles that lacked ambiguous nucleotides (Ns) and searched for sequence contexts specific to each enzyme. Specifically, we identified GCH sites (for M.CviPI), adenines (for Hia5/M.EcoGII), and cytosines (for DddB/SsDddA) on both DNA strands to estimate their densities.

We chose to search both strands because third-generation sequencing technologies (e.g., PacBio) can detect enzymatic marks on both strands simultaneously. In contrast, short-read bisulfite sequencing datasets (e.g., NOMe-seq sequenced with Illumina) provide information from only one strand; therefore, the estimated densities should be halved when applied to such data.

### NOMe-seq experiment for mouse ES cells

#### Design and validation of PCR primers for amplicons

NOMe-seq primers were designed against *in-silico* bisulfite converted DNA sequences using Primer3 with minor modifications (21). Targeted genomic sequences for primer hybridization lacked CpG and GpC dinucleotides. Amplicon sizes ranged from 300 bp to 479 bp and the annealing temperatures between 50°C and 60°C. Primers were commercially synthesized and the amplification products were validated by agarose gel electrophoresis (Supplementary Table 1).

#### NOMe-seq experiment

Growing JM8 mESCs were trypsinized and washed once with cold phosphate buffered saline (PBS). 250.000 mESC were permeabilized with 1ml ice-cold lysis buffer (10 mM Tris pH 7.4, 10 mM NaCl, 3 mM MgCl2, 0.1 mM EDTA, 0.5% Igepal) for 10 min on ice. Cells were spun for 5 min at 600g and 4°C and the pellet was resuspended in 250 µl cold wash buffer (10 mM Tris pH 7.4, 10 mM NaCl, 3 mM MgCl2, 0.1 mM EDTA). After centrifugation for 5 min at 600g and 4°C, the pellet was resuspended in 94.5 µl of 1x GC Reaction Buffer (M0227L, NEB). Footprinting was performed by the addition of 200U M.CviPI (M0227L, NEB) in a final concentration of 0.65 mM SAM (M0227L, NEB), 300 mM sucrose and 1xGC Reaction Buffer.

After 7.5 minutes at 37°C, the reaction was supplemented with 100U of M.CviPI and SAM concentration was increased to 1 mM. Nuclei were further incubated for 7.5 minutes at 37°C before stopping the reaction with pre-warmed stop buffer (10 mM Tris pH 7.4, 200 mM NaCl, 5 mM EDTA, 0.5% SDS and 200 µg/ml Proteinase K). After 16h incubation at 55°C and 1200 rpm, samples were RNAse treated for 30 min at 37°C. DNA was purified with the addition of 1:1 v/v Phenol:Chloroform:Isoamyl Alcohol solution (15593031, ThermoFisher), mixed and spun for 10 min at maximum speed. The supernatant was transferred to a new DNA LoBind microcentrifuge tube (Eppendorf) and further purified by adding 1:1 v/v chloroform. After mixing and spinning for 10 min at maximum speed, the supernatant was transferred to a new tube and supplemented with 1µl of Glycogen, 1:2 v/v ethanol 100% and 1:0.1 v/v 3M sodium citrated. DNA was precipitated at −80°C for 1h. Samples were centrifuged for 30 min at maximum speed and 4°C. The supernatant was discarded, the pellet was washed with 500 µl cold 70% ethanol and spun for 10 min at maximum speed and 4°C. DNA pellets were allowed to dry and were resuspended in nuclease-free water and quantified with Qubit 1x dsDNA High Sensitivity kit (Invitrogen). Around 1.2 µg of DNA were bisulfite converted with EpiTect Bisulfite Kit (59104, QIAGEN) according to manufacturer’s instructions. Amplicons were individually amplified in a 96-well plate using AmpliTaq Gold DNA polymerase (N8080249, Applied biosystems) in a total volume of 25 µl with 0.4uM primers and ∼12ng DNA as measured prior to bisulfite conversion (95°C, 9 min; 20 touchdown cycles [95°C, 30 s; 56 to 52°C, 30 s; 72°C, 1 min]; 36 cycles [95°C, 30 s; 52°C, 30 seconds; 72°C, 1 min]; 5min at 72°C). 5µl of each PCR amplified amplicon were pooled, purified with 1X AMPureXP beads (A63881, Beckman Coulter) and quantified using Qubit 1x dsDNA High Sensitivity kit. Libraries were prepared with NEBNext ChIP-seq Library Prep (NEB, E6240) according to manufacturer’s instructions. Quality control of the libraries was performed with Fragment Analyzer (Agilent). Libraries were sequenced 250 x 250 bp on an Illumina MiSeq instrument.

#### Preprocessing and alignment

Paired-end NOMe-seq raw sequencing reads were preprocessed using *Trim Galore* (69) to remove Illumina adapters and low-quality bases (parameters: --paired --illumina --quality 20 --length 125 --clip_R1 5 --clip_R2 5). Trimmed reads were aligned to the amplicon sequences using the R package *QuasR* (70) with the *qAlign* function and the following parameters: paired = “fr”, bisulfite = “undir”, and alignmentParameter = ‘-X 700 -k 2 --best --strata’.

#### Data filtering and fetching

Methylation calling and data retrieval from BAM files were performed using *fetchNOMe*, an R package developed for this study. Briefly, cytosine methylation calling was conducted on individual sequenced fragments (i.e., single DNA molecules) by analyzing C-to-T and G-to-A substitutions resulting from bisulfite conversion.

The following sequence contexts were considered:

- Accessibility information: Cytosines within the GCH context (GCA/GCC/GCT)
- Endogenous CpG methylation: Cytosines within the WCG context (ACG/TCG)
- Bisulfite conversion efficiency: Cytosines within “bisC” contexts (ACA/ACC/ACT/TCA/TCC/TCT) were used to identify fragments with incomplete bisulfite conversion. Fragments showing >10% “bisC” methylation were considered incompletely converted and were filtered out. A minimum of 10 “bisC” positions was required to evaluate bisulfite conversion efficiency.

Co-occurrence count tables (Figure 1D), required for footprint spectral analysis, were generated for each amplicon using the *get_ctable_from_bams* function. Spacings up to 200 bp were considered, and protected positions at both ends of each molecule were removed to minimize artifacts caused by truncated footprints at the amplicon edges.

Data matrices containing accessibility information for each molecule within an amplicon were generated using the *get_data_matrix_from_bams* function in the *fetchNOMe* package.

#### Classification of the amplicons

We classified amplicons using publicly available ChIP-seq datasets for TFs Ctcf and Rest (41) and chromatin marks (19). DNAme levels for each amplicon were estimated using WCG methylation from the NOMe-seq experiment conducted for this study.

#### Footprint spectral analysis

Footprint spectral analysis was performed on co-occurrence count tables for each amplicon using the *infer_footprints_vb* function from the *nomeR* package, with the parameter *ftp_bg_model* set to *“informative_prior”* and footprint lengths ranging from 10 to 200 bp. Non-informative Dirichlet distributions were used as priors for footprint coverages, while Beta distributions served as priors for emission probabilities with parameters: bg_protect_mean = 0.05, bg_protect_totcount = 40, ftp_protect_mean = 0.95, and ftp_protect_totcount = 40.

The means and standard deviations of the inferred parameters were calculated using 4,000 samples drawn from the mean-field variational approximation of the posterior distribution.

#### Predicting nucleosome footprints

To predict nucleosome footprints, we selected amplicons that lacked TF footprints and calculated posterior probabilities for footprints ranging from 100 to 150 bp using the *predict_footprints* function from the *nomeR* package. Based on the footprint spectral analysis, we set the following parameters:

- Expected background coverage: 0.54
- Emission probabilities (probability of observing protected positions):

- Background: 0.01
- Footprints: 0.92

To avoid bias towards specific footprint lengths, we parameterized the prediction such that prior start probabilities were equal for all footprints between 100 and 150 bp.

After calculating posterior probabilities for each footprint, we summed them to identify molecules containing footprints with total start probabilities above 0.05. Finally, we centered all molecules relative to the start sites of the most confidently predicted nucleosome footprints (i.e., those with the highest start probability) (Figure 4D).

### Analysis of publicly available genome-wide NOMe-seq data for mESC

#### Data acquisition, preprocessing and alignment

Previously published (29) paired-end genome-wide bulk NOMe-seq datasets for mESCs were downloaded from the GEO repository (GSE78140, samples GSM2200651, GSM2200652, GSM2200653 and GSM2200654). Illumina adapter trimming was performed using Trim Galore with parameters “--clip_R1 15 --clip_R2 15 --trim1 --paired.” Reads were aligned to the mouse genome (mm10) using Bismark (v0.22.3) (71) with the parameters “--bowtie2 --local - X 2000 --non_directional.”

#### Genome-wide footprint spectral analysis

Co-occurrence count tables were generated from BAM files using the *get_ctable_from_bams* function from the *fetchNOMe* R package for 10 kb genomic tiles, excluding tiles overlapping ENCODE blacklisted regions. Only fragments longer than 50 bp with informative position densities above 7% were included (parameters: min_frag_data_len = 50, min_frag_data_dens = 0.07).

For spectral analysis, count tables were aggregated separately for autosomes/sex chromosomes (chr1–19, chrX, chrY), and mitochondrial DNA (chrM). Bayesian parameter inference was conducted using the *infer_footprints_vb* function from the *nomeR* package with the *ftp_bg_model* parameter set to *“informative_prior”* and footprint lengths ranging from 5– 200 bp. Non-informative Dirichlet distributions were used as priors for footprint coverages, while Beta distributions served as priors for emission probabilities with parameters: bg_protect_mean = 0.05, bg_protect_totcount = 40, ftp_protect_mean = 0.95, and ftp_protect_totcount = 40. Parameter means and standard deviations were estimated from 4,000 samples drawn from the mean-field variational approximation of the posterior distribution.

#### Predicting nucleosome footprints

To detect nucleosome footprints, 100,000 random 1 kb tiles were selected, and accessibility data were extracted using *get_data_matrix_from_bams* from the *fetchNOMe* R package. Posterior probabilities for footprints (100–150 bp) were calculated using *predict_footprints* from *nomeR*, with background coverage set to 0.51, background emission probability to 0.01, and footprint emission probability to 0.87. Equal prior start probabilities were assigned to all footprint lengths to prevent bias. Molecules with maximum (across all positions) total (sum across all footprints) start probabilities above 0.05 were identified, and centered relative to the most confident footprint starts (Figure 4F).

#### Analysis of ChIP-seq datasets for TFs

Analysis of ChIP-seq datasets and binding sites for TFs Ctcf and Rest related to Figures 4A-4B, 5A-B was described previously (41). Briefly, ChIP-seq datasets that are publicly available at the GEO repository (GSE112136) were aligned to the mouse genome using QuasR (70) with default parameters. Position frequency matrices from JASPAR2016 (39) were used to obtain position weight matrices (PWMs) for Ctcf and Rest. Genome-wide binding sites were predicted with the *matchPWM* function from *Biostrings* (v2.46.0) using a minimum score threshold of 10.0. BigWig files for genomic track visualization (Figures 5D, S3C) were generated with *qExportWig* from *QuasR* (*binsize=50, scaling=1e6*).

Analysis of ChIP-seq datasets and *de novo* motif finding related to Figure 5C-5D, Extended Data Figure 2, were described previously (42). Briefly, publicly available ChIP-seq datasets for TFs were downloaded from the GEO repository: Ctcf (GSM747535 (48)), Esrrb (GSM2417188 (43)), Klf4 (GSM2417144 (43)), MafK (GSM1003809 (49)), Nanog (GSM2417187 (43)), Nrf1 (GSM1891642 (4)), Oct4 (GSM2417142 (43)), p53 (GSM647224 (47)), Prdm14 (GSM623989 (44)), Rest (GSM671095 (45)), Sox2 (GSM2417143 (43)), Tcfcp2I1 (GSM288350 (46)), Zc3h11a (GSM1003810 (49)), Zfx (GSM288352 (46)). ChIP-seq datasets were aligned with QuasR (70) followed by peak calling with MACS2 (72) and *de novo* motif finding was performed for the top 500 peaks using the function *findMotifsGenome.pl* from the HOMER toolset (73) (curated motifs are displayed in Figure 5C, Extended Data Figure 2C). Genomic locations for TF motifs were determined with the *matchPWM* function from the *Biostrings* R package using resulting positional weight matrices for TF motifs and selecting matches with log2(odds) score above 10. Finaly, log2 ChIP-seq enrichments were quantified at 201 bp regions centered around TF motif modpoints as log2 ratios of ChIP CPMs and CPMs for corresponding Inputs. (Extended Data Figure 2A).

#### Footprint spectral analysis for TFs

For the spectral analysis of NOMe-seq data (29) at TF binding sites we selected motifs with log2 enrichments above 1 and containing at least 1 GCH within 30bp around motif centers. Next, we investigated co-occupancy of TFs by calculating pairwise Jaccard indexes as ratios of genomic lengths for intersections and unions of the 100bp regions centered around motifs (Extended Data Figure 2B). For TFs that showed co-occupancy, i.e. Nanog-Oct4-Sox2 and Ctcf-Zc3h11a, we subdivided TF regions into the following groups (Extended Data Figure 2B):

1. “Nanog-Sox2-Oct4” contains 100bp regions where all 3 TFs co-occupy
2. “Nanog” contains 100bp regions where binding of only Nanog was observed
3. “Sox2” contains regions where binding of only Sox2 was observed
4. “Oct4” contains regions where binding of only Oct4 was observed.

Similarly,

1. “Ctcf-Zc3h11a” contains regions where binding of both Ctcf and Zc3h11a was observed
2. “Ctcf” contains regions where only binding of Ctcf was observed
3. “Zc3h11a” contains regions where only binding of Zc3h11a was observed.

Next we collected accessibility information from NOMe-seq dataset (29) for molecules overlapping selected regions and calculated posterior probabilities for TF footprints (15-80bp) and nucleosomes (100-150bp) using the function *predict_footprints* assuming coverages for background, nucleosomes and TFs to be 0.80, 0.15 and 0.05 respectively. We also set emission probabilities for background and footprints to be 0.02 and 0.85 respectively.

We calculated BG and TF scores by applying log-ratio transformations (74–76) to predicted coverages as follows:

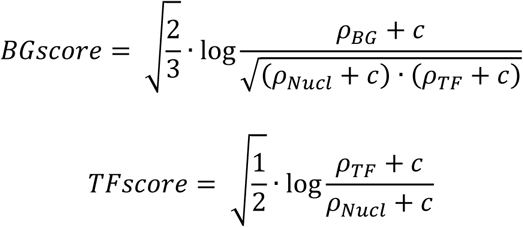

where 𝜌*_BG_*, 𝜌*_Nucl_* and 𝜌*_TF_* are calculated posterior coverage probabilities and 𝐵 = 0.1 is a pseudocount.

Next, we calculated average TF scores across positions covered by TF motifs and standardized them by subtracting the mean and dividing by the standard deviation (Z-score) across all motifs included into the analysis. We ordered molecules by standardized TF scores and chose a cutoff of 0.4 to identify molecules that are likely to contain footprints for TFs. We illustrated the raw accessibility of GCHs in form of heatmaps (Figure 5C, Extended Data Figure 2C) where rows correspond to single molecules and columns correspond to relative position within 101bp regions centered around motif midpoints.

Finally, for the footprint spectral analysis we removed molecules with non-positive standardized TF scores, selected molecules with standardized TF scores above 0.4 but not fewer than 10,000, collected co-occurrence count tables for 101bp regions around motif centers and performed spectral analysis using the function *infer_footprints_vb* for footprint lengths ranging from 10 to 200bp (Figure 5D, Extended Data Figure 2D).

#### Analysis of ATAC-seq dataset

Paired-end ATAC-seq data (53) were downloaded from GEO (GSE85632; GSM2279944, GSM2279945). Nextera adapters were removed using Trim Galore (69) with parameters “-- nextera --stringency 5” and trimmed reads were aligned to mouse genome using STAR (77) with parameters “--alignIntronMin 1 --alignIntronMax 1 --alignEndsType EndToEnd -- alignMatesGapMax 1000 --outFilterMatchNminOverLread 0.85”. Prior to peak calling, biological replicates were merged and the resulting BAM file was converted to a BED file using the bamToBed utility from the bedtools toolset (78). Peak calling was performed using MACS2 with parameters “-g mm --keep-dup all -q 0.10 --nomodel --shift -100 --extsize 200”. Bigwig files for visualization (Figures 6A, Extended Data Figure 3D) were generated using the function *qExportWig* from the *QuasR* package with parameters “binsize=100L, scaling=1e+6, mapqMin=255L, mapqMax=255L, createBigWig=TRUE, useRead=“any”, pairedAsSingle=TRUE, absIsizeMax = 1000”.

#### De novo identification of open chromatin regions and regions with TF binding

For the proof-of-principle we selected 11 genomic loci (length 1-1.5Mb) characterized by higher densities of ATAC-seq peaks (53) and binding sites for TFs Ctcf and Rest (41) in mESCs. For the chosen loci we collected GCH accessibility information for all molecules longer than 50bp with densities of information positions above 5% from the genome-wide NOMe-seq dataset (29) using the function *get_molecule_data_list_from_bams* (*fetchNOMe*).

For GCH accessibility data in each molecule we calculated posterior coverage probabilities of footprints for TFs (15-80bp) and nucleosomes (100-150bp) assuming coverages for background, nucleosomes and TFs to be 0.45, 0.50 and 0.05 respectively. We also set emission probabilities for background and footprints to be 0.01 and 0.90 respectively. Given resulting posterior coverage probabilities we calculated BG and TF scores by applying log-ratio transformation as described above.

Based on BG and TF scores we identified GCHs that are likely to be accessible (covered by background model) or covered by a TF footprint within each molecule using the following rules: if the predicted BG score for a GCH is above 0.5 we identify it as a background (BG), otherwise if the predicted TF score for a GCH is above 0.5 and BG score is below 0.5 we identify it as covered by the TF footprint (TF).

For sliding windows of 500bp with step 250bp we counted numbers of GCHs identified as BG and TF across all molecules and performed a one-tailed binomial test using the R function *binom.test* for the null-hypothesis that the fraction of BG or TF for each particular window equals the expected fraction across all molecules (total number of BG or TF GCHs divided by total number of GCHs) with alternative hypothesis that the fraction is greater than expected. The p-values were corrected for multiple testing using Benjamini-Hochberg method with R function *p.adjust*. The resulting fractions of BG and TF GCHs and −log10 transformed False Discovery Rates (FDR) for every window are displayed as volcano plots in Extended Data Figure 3A-3B as well as genomic tracks “occupancy” and “stat.sign.” in Figure 6A, Extended Data Figure 3D.

Endogenous DNA methylation levels were estimated from WCG sites using *get_molecule_data_list_from_bams* (parameter whichContext = “WCG”). WCG Methylation levels were calculated as the fraction of methylated WCGs among covered sites, excluding sites with <5 reads.

Methylation levels for sliding windows were calculated as average across all WCG overlapping corresponding windows and displayed as genomic tracks “DNAme” in Figures 6A, Extended Data Figure 3D, define point colors in volcano plots in Extended Data Figure 3A-3B as well as displayed as violin plots in Figure 6C.

Classification of sliding windows with respect to statistically significant enrichment of TF footprints and overlap with ATAC-seq summits were done as follows. Windows with FDR ≤ 0.01 for enrichment of TF footprints according to analysis of NOMe-seq data were considered as statistically significant. ATAC-seq summits with q-values ≥ 2 were selected, overlapped with sliding windows, the following grouping was applied: windows with statistically significant enrichment of TF footprints overlapping or not overlapping ATAC-seq summits were assigned groups “ATAC & NOMe” and “NOMe only” respectively, whereas windows overlapping ATAC-seq summits and lacking statistically significant enrichment of TF footprints were assigned a group “ATAC only” (Figure 6B). All other windows were classified as “other” (Figure 6C, Extended Data Figure 3C).

Classification of sliding windows with respect to overlap with confirmed ChIP-seq summits annotated in REMAP2022 database (54) was done as follows. BED file with all annotated peaks for mESC was downloaded from https://remap.univ-amu.fr/storage/remap2022/mm10/MACS2/BIOTYPE/mESC/remap2022_mESC_all_macs2_mm10_v1_0.bed.gz. Samples names in the BED file were parsed and only ChIP-seq experiments for WT mESC and peaks/summits with score ≥ 10 were retained for further analysis. The sliding windows were overlapped with resulting ChIP-seq summits and categorized as “with” or “without” REMAP (Figure 6B-6C, Extended Data Figure 3C). Genomic tracks “REMAP2022” for Figure 6A, Extended Data Figure 3D were obtained by calculating coverage of selected REMAP2022 peaks within the investigated regions.

### Software availability

The R package *fetchNOMe*, developed for fast retrieval of SMF data and extraction of statistics required for footprint spectral analysis from BAM files generated by NOMe-seq experiments, is available on GitHub at https://github.com/fmi-basel/gpeters-fetchNOMe.

The R package *nomeR*, designed for performing footprint spectral analysis and predicting footprint positions within individual SMF molecules, is available on GitHub at https://github.com/fmi-basel/compbio-nomeR.

## AUTHORS CONTRIBUTIONS

E.A.O conceived the statistical model, developed R packages *fetchNOMe* and *nomeR*, designed and performed the *in-silico* simulations and interpreted the data. L.G.-T. designed and performed amplicon-based NOMe-seq experiment in mESCs. E.A.O and L.G-T. analyzed and interpreted the data from the amplicon-based NOMe-seq experiment. E.A.O designed and performed the analysis of publicly available genome-wide NOMe-seq dataset for mESCs. A.H.F.M.P. conceived and supervised the overall project. E.A.O. wrote the manuscript with input from all authors.

## ACKNOWLEDGEMENTS

We gratefully acknowledge M. Stadler and L. Burger for providing annotations for TF binding sites. We also thank members of the Peters laboratory and FMI colleagues, especially M. Gill, G. Fanourgakis, M. Stadler, C. Soneson, L. Burger and M. Schwaiger, for critical reading of the manuscript. This research was supported by the Novartis Research Foundation, the Swiss National Science Foundation (31003A-172873) and the European Research Council (ERC) under the European Union’s Horizon 2020 research and innovation program (grant agreement ERC-AdG 695288 – Totipotency).

