## Supplementary Text for "A statistical model for quantitative analysis of single-molecule footprinting data"

### Introduction

Regulation of gene expression by DNA binding proteins and complexes, such as sequence specific transcription factors (TFs), is a key step that drives a plethora of biological processes. During the past decade, remarkable achievements have been made using ChIP-seq and ATAC-seq assays to understand how TFs regulate their targets. However, these assays provide only averaged information of larger cell populations. Hence, such assays are unable to detect states of binding at single-DNA-molecule resolution. In contrast, emerging single-molecule footprinting (SMF) technologies, such as the Nucleosome Occupancy and Methylome Sequencing (NOMe-seq) assay, provide information about states of binding sites at single-DNA-molecule resolution and allow better molecular characterization of inherently stochastic processes of protein-DNA interactions.

Here, we develop a statistical framework for analyzing datasets generated by SMF assays. First, we introduce a stochastic process that we use for modeling the data. Second, we demonstrate how parameters, such as abundances of footprints and noise levels, can be inferred from the data. Finally, we introduce an approach for disentangling footprints of different sizes by predicting their positions within each single molecule, which can be used for rigorous quantitative analysis of SMF datasets.

### Stochastic process

Single-molecule footprinting (SMF) assays generate data that provide information about accessibility of substrate nucleotides for enzymatic marking by a chosen exogenous enzyme, e.g. GpC when M.CviPI (Kelly et al., 2012), adenines when m6A-Mtase such as Hia5 (Stergachis et al., 2020) or M.EcoGII (Abdulhay et al., 2020; Murray et al., 2018), or cytosines when deaminase such as DddB (He et al., 2024) or SsDddA (Swanson et al., 2024) are chosen respectively.

For development of our statistical framework for single-molecule footprinting data we utilize a model previously described for biophysical modeling of protein-DNA interactions (McGhee and von Hippel,

1974; Ozonov and Nimwegen, 2013; Raveh-Sadka et al., 2009; Teif and Rippe, 2010) and prediction of regulatory sequences (van Nimwegen, 2007).

We assume that single-molecule footprinting data are generated by a stochastic process that randomly places $K$ footprints of lengths $\boldsymbol{l} = (l_1, \dots, l_K)$ with certain prior start probabilities $\boldsymbol{\pi} = (\pi_1, \dots, \pi_K)$ in a sequence of length $L$, where index $1$ is reserved for accessible position (i.e. background), therefore $l_1 = 1$ and $\pi_1$ represents prior probability for placing an accessible position during the described stochastic process. Also, as vector $\boldsymbol{\pi}$ represents probabilities that a position is a start of a footprint, it is a point on a K-dimensional simplex, i.e. $\sum_{i=1}^{K} \pi_i = 1$.

We assume that footprints cannot overlap and do not interact with each other, and footprints emit 0 (unprotected or methylated position) and 1 (protected or unmethylated position) with certain emission probabilities, which constitute observed data acquired from a SMF experiment (Figure S1).

To calculate probability that a certain footprint $\omega$ occupies a stretch of a sequence starting at position $n$ (Figure S1) we consider all possible configurations of footprints on a sequence and for each configuration we assign a statistical weight $W$, that depends on the underlying sequence, footprint and background emission probabilities as well as on footprint's lengths and prior start prior probabilities.

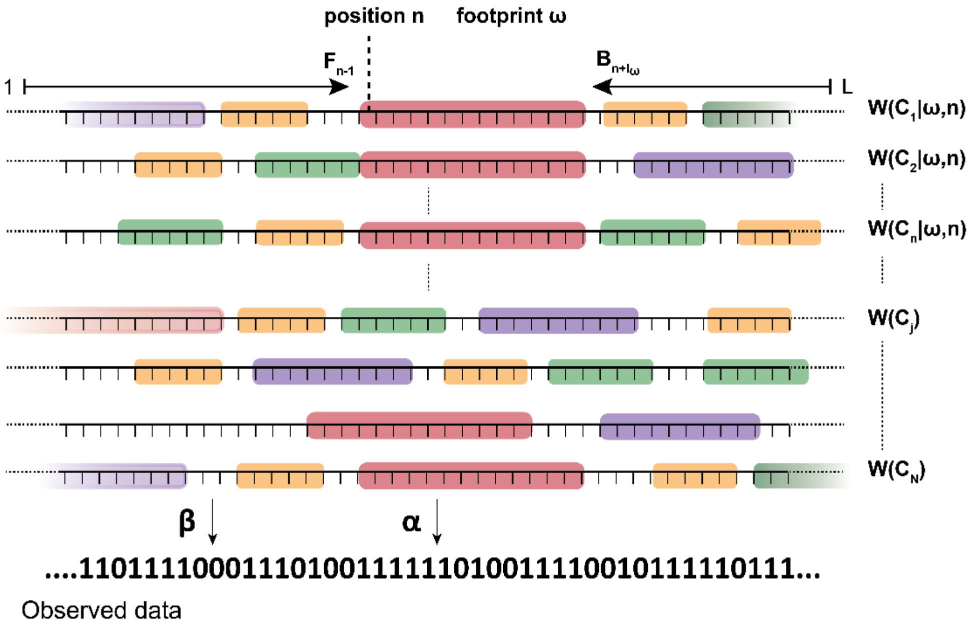


Figure.S7¡.Illustration.of.possible.configurations.of..K.footprints.on.a.sequence.with.length.L¡.Top. configurations.represent.a.set..$\boldsymbol{C}(\boldsymbol{w}, \boldsymbol{n})$.which.contain.all.configurations.with.a.footprint.$\omega$.at.position.$n$. with.statistical.weights.$W(C_t|w,n)$¡.Example.of.observed.data.(for.simplicity.data.for.all.positions.are. available).is.shown.at.the.bottom¡

Next, to calculate that a certain footprint $\omega$ starts from position $n$ we sum all statistical weights which contain the footprint $\omega$ at position $n$ and normalize it to total sum of statistical weights across all possible configurations.

$$P(n|\omega, \boldsymbol{\pi}) = \frac{\sum_{C_t \in \boldsymbol{C}(\boldsymbol{w},\boldsymbol{n})} W(C_t|w,n)}{\sum_{C_j} W(C_j)},$$

where $\boldsymbol{C}(\boldsymbol{w}, \boldsymbol{n})$ is a set of configurations with footprint $\omega$ starting at position $n$ (Figure S1).

The sums of statistical weights can be efficiently calculated using so-called transfer-matrix of dynamic-programming techniques (van Nimwegen, 2007), with the recursive relationships and boundary conditions defined below

$$F_n = \sum_{\omega=1}^{K} \pi_\omega P\big(S_{n-l_\omega+1}\big|\omega\big) F_{n-l_\omega} \tag{1}$$

$$B_n = \sum_{\omega=1}^{K} \pi_\omega P(S_n|\omega) B_{n+l_\omega} \tag{2}$$

$$F_0 = 1;\ B_{L+1} = 1 \tag{3}$$

$$P_{start}(n|\omega, \boldsymbol{\pi}) = \frac{F_{n-1}\pi_\omega P(S_n|\omega) B_{n+l_\omega}}{F_L} \tag{4}$$

$$P_{end}(n|\omega, \boldsymbol{\pi}) = P_{start}(n - l_\omega + 1|\omega, \boldsymbol{\pi}) = \frac{F_{n-l_\omega}\pi_\omega P\big(S_{n-l_\omega+1}\big|\omega\big) B_{n+1}}{F_L} \tag{5}$$

Where $P_{start}(n|\omega, \boldsymbol{\pi})$ and $P_{end}(n|\omega, \boldsymbol{\pi})$ are **posterior** probabilities that footprint $\omega$ starts or ends at position $n$ and $P(S_n|\omega)$ is a **prior** probability that footprint $\omega$ covers a region starting from $n$ until $n + l_\omega - 1$, which introduces sequence or positional specificity for footprints.

Note that in order to calculate the probability that a position $n$ is covered by a footprint $\omega$ we sum starting probabilities for all positions where footprint $\omega$ overlaps position $n$, i.e.

$$\rho(n|\omega, \boldsymbol{\pi}) = \sum_{i=n-l_\omega+1}^{n} P(i|\omega, \boldsymbol{\pi}) \tag{6}$$

Also, note that the sum of coverage posterior probabilities across all footprints, including background, must equal 1, i.e.

$$\sum_{\omega=1}^{K} \rho(n|\omega, \boldsymbol{\pi}) = 1 \tag{7}$$

which is equivalent to the condition that the total probability of a position $n$ being covered by anything, including background, must equal 1. On the other hand, the sum of posterior probabilities to start or end at position $n$ across all footprints does not necessarily equal 1.

### Model reformulation

Here we apply reformulation of the model used in the code of (Arnold et al., 2012; Ozonov and Nimwegen, 2013) which improves numerical stability as well as helps us to derive some important relationships later in this document.

Our objective is to derive recursive relationships (1) and (2) and posterior probabilities (4) and (5) in terms of

$$R_n = \frac{F_n}{F_{n-1}};\ R_n^{(B)} = \frac{B_n}{B_{n+1}};\ X_n = \frac{1}{R_n};\ X_n^{(B)} = \frac{1}{R_n^{(B)}} \tag{8}$$

By definition

$$\begin{aligned} F_n &= R_n F_{n-1} = R_n R_{n-1} F_{n-2} = \cdots = \prod_{i=1}^{n} R_i\,; \\ B_n &= R_n^{(B)} B_{n+1} = R_n^{(B)} R_{n+1}^{(B)} B_{n+2} = \cdots = \prod_{i=n}^{L} R_i^{(B)} \end{aligned} \tag{9}$$

Then, substituting $F_n$ and $B_n$ in (1) and (2) using (8) and (9)

$$R_n = \sum_{\omega} \left[ \pi_\omega P\left(S_{n-l_\omega+1} \middle| \omega\right) \prod_{i=n-l_\omega+1}^{n-1} X_i \right];$$

$$R_n^{(B)} = \sum_{\omega} \left[ \pi_\omega P(S_n|\omega) \prod_{i=n+1}^{n+l_\omega-1} X_i^{(B)} \right]$$

or

$$\begin{aligned} X_n &= \frac{1}{\sum_\omega \left[ \pi_\omega P\left(S_{n-l_\omega+1} \middle| \omega\right) \prod_{i=n-l_\omega+1}^{n-1} X_i \right]}; \\ X_n^{(B)} &= \frac{1}{\sum_\omega \left[ \pi_\omega P(S_n|\omega) \prod_{i=n+1}^{n+l_\omega-1} X_i^{(B)} \right]} \end{aligned} \tag{10}$$

Then, posterior probabilities (4) and (5) are

$$P_{start}(n|\omega, \boldsymbol{\pi}) = \pi_\omega P(S_n|\omega) \cdot \prod_{i=n}^{n+l_\omega} X_i \cdot Z_{n+l_\omega+1} \tag{11}$$

$$P_{end}(n|\omega, \boldsymbol{\pi}) = \pi_\omega P\left(S_{n-l_\omega+1}\middle|\omega\right) \cdot \prod_{i=n-l_\omega+1}^{n} X_i \cdot Z_{n+1} \tag{12}$$

where

$$Z_n = \prod_{i=n}^{L} \frac{X_i}{X_i^{(B)}} = \frac{X_n}{X_n^{(B)}} \cdot Z_{n+1} \tag{13}$$

with boundary conditions

$$\begin{gathered} X_0 = 1; \\ X_{L+1}^{(B)} = 1; \\ Z_{L+1} = 1 \end{gathered} \tag{14}$$

### Boundary conditions for infinite sequence

Let's consider a sequence of length $L$ bp, which is a part of much longer sequence. In other words, the sequence of interest for which we would like to calculate posterior probabilities is a part of infinite

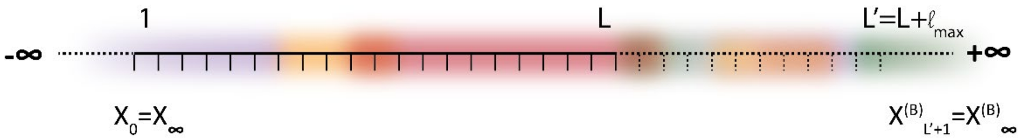

Figure.S8¡.Illustration.for.a.sequence.of.length.L.being.a.part.of.an.infinite.sequence¡.All.positions. outside.of.[7·L].have.no.positional.or.sequence.specificities.for.footprints¡

sequence (Figure S2).

In reality, our sequence of interest would be a sequenced single molecule, which usually is about 500-600 bp, in the context of a chromosome that is many orders of magnitude longer.

We assume that anywhere outside the region of interest [1;L] footprints do not have any positional specificity, i.e. $P(S_n|\omega) = 1 \forall \omega, n \notin [1; L]$. Our objective is, therefore, to calculate posteriors (11) or (12), but instead of boundary conditions (14) we would like to use boundary conditions $X_\infty = \lim_{a \to -\infty} X_{[a;0]}, X_\infty^{(B)} = \lim_{b \to \infty} X_{[L+1;b]}^{(B)}$ which would take into account the assumption that our sequence of interest is a part of an infinite sequence.

Intuitively, both $X_\infty$ and $X_\infty^{(B)}$ equal 1, as by definition they are both ratios of neighboring forward and backward partition sums (see (8)). Indeed, substituting $X_n = X_\infty$ in the recursive relationships (10) we can see that $X_\infty$ equals to a root of polynomial $\sum_\omega \pi_\omega X_\infty^{l_\omega} - 1 = 0$, which has a single positive root $X_\infty = 1$ when $\sum_{\omega=1}^{K} \pi_\omega = 1$.

For calculation of boundary condition for $Z_n$ we can use normalization (7), i.e. the requirement that the total probability that a position is covered by any footprint, including background, must equal 1. Let $l_{max}$ be the length of the longest footprint. Also, we consider a region extended by $l_{max}$ with the length $L' =$

$L + l_{max}$, and as before footprints lack sequence or positional specificity outside of [1;L], i.e. $P(S_n|\omega) = 1 \forall \omega, n \notin [1; L]$. Then, our objective is to find a boundary condition $Z_{L'+1}$ that fulfils the normalization requirement, i.e. $\sum_{\omega=1}^{K} \rho(L'|\omega, \boldsymbol{\pi}) = 1$.

Using (12) that probability that position $L'$ is covered by a footprint $\omega$ is

$$\rho(L'|\omega, \boldsymbol{\pi}) = \sum_{n=L'}^{L'+l_\omega-1} P_{end}(n|\omega) = \pi_\omega \sum_{n=L'}^{L'+l_\omega-1} Z_{n+1} \prod_{i=n-l_\omega+1} X_i \quad (15)$$

From equation (13) it follows that

$$Z_n = Z_{n-1} \cdot \frac{X_{n-1}^{(B)}}{X_{n-1}} = Z_{n-2} \cdot \frac{X_{n-2}^{(B)}}{X_{n-2}} \cdot \frac{X_{n-1}^{(B)}}{X_{n-1}} = \cdots = Z_{n-d} \cdot \prod_{i=n-d}^{n-1} \frac{X_i^{(B)}}{X_i} \quad (16)$$

In particular,

$$Z_{n+1} = Z_{L'+1} \cdot \prod_{i=L'+1}^{n} \frac{X_i^{(B)}}{X_i} \quad (17)$$

We can express (15) in terms of $Z_{L'+1}$ using (17).

$$\begin{aligned}
\rho(L'|\omega, \boldsymbol{\pi}) &= \pi_\omega \left[ Z_{L'+1} \prod_{i=L'-l_\omega+1}^{L'} X_i + \sum_{n=L'+1}^{L'+l_\omega-1} Z_{n+1} \prod_{i=n-l_\omega+1}^{n} X_i \right] \\
&= \pi_\omega \left[ Z_{L'+1} \cdot \prod_{i=L'-l_\omega+1}^{L'} X_i + Z_{L'+1} \cdot \sum_{n=L'+1}^{L'+l_\omega-1} \prod_{i=L'+1}^{n} \frac{X_i^{(B)}}{X_i} \cdot \prod_{i=n-l_\omega+1}^{n} X_i \right] \\
&= \pi_\omega \cdot Z_{L'+1} \cdot \left[ \prod_{i=L'-l_\omega+1}^{L'} X_i + \sum_{n=L'+1}^{L'+l_\omega-1} \prod_{i=n-l_\omega+1}^{L'} X_i \cdot \prod_{i=L'+1}^{n} \frac{X_i^{(B)}}{X_i} \cdot X_i \right] \\
&= \pi_\omega \cdot Z_{L'+1} \cdot \left[ \prod_{i=L'-l_\omega+1}^{L'} X_i + \sum_{n=L'+1}^{L'+l_\omega-1} \prod_{i=n-l_\omega+1}^{L'} X_i \cdot \prod_{i=L'+1}^{n} X_i^{(B)} \right]
\end{aligned}$$

Since $X_i^{(B)} = X_\infty^{(B)} = 1, \forall i > L$

$$\rho(L'|\omega,\boldsymbol{\pi}) = \pi_\omega \cdot Z_{L'+1} \cdot \left[ \prod_{i=L'-l_\omega+1}^{L'} X_i + \sum_{n=L'+1}^{L'+l_\omega-1} \prod_{i=n-l_\omega+1}^{L'} X_i \right]$$

$$= \pi_\omega \cdot Z_{L'+1} \cdot \sum_{n=L'}^{L'+l_\omega-1} \prod_{i=n-l_\omega+1}^{L'} X_i \quad (18)$$

Therefore, from (18) and the normalization $\sum_{\omega=1}^{K} \rho(L'|\omega,\boldsymbol{\pi}) = 1$, we get the following boundary conditions for recursive relationships (10) and (13).

$$Z_{L'+1} = \frac{1}{\sum_\omega \pi_\omega \cdot \sum_{n=L'}^{L'+l_\omega-1} \prod_{i=n-l_\omega+1}^{L'} X_i}$$

$$X_0 = X_\infty = 1; \quad (19)$$

$$X_{L'+1}^{(B)} = X_\infty^{(B)} = 1;$$

Importantly, using boundary conditions for recursive relationships for an infinite sequence allows us to avoid boundary effects, such as statistical positioning (Kornberg and Stryer, 1988; Raveh-Sadka et al., 2009) that may distort predicted posterior probabilities, given that sequenced fragments are only 500-600bp long.

Interestingly, if we assume no sequence specificity across the entire infinite sequence, i.e. $P(S_n|\omega) = 1 \; \forall \; \omega, n$ we get from equations (11) and (19) very intuitive relationships between prior probabilities $\boldsymbol{\pi}$ in the described stochastic process and posterior probabilities. Indeed, when $P(S_n|\omega) = 1 \; \forall \; \omega, n$ is assumed, it follows that $X_n = X_n^{(B)} = 1$ and $Z_n = \frac{1}{\sum_{\omega=1}^{K} \pi_\omega l_\omega}, \forall n$. Therefore,

$$P_{start}(\omega,\boldsymbol{\pi}) = P_{end}(\omega,\boldsymbol{\pi}) = \frac{\pi_\omega}{\sum_{\omega'} \pi_{\omega'} l_{\omega'}} \quad (20)$$

$$\rho_\omega \equiv \rho(\omega,\boldsymbol{\pi}) = \frac{\pi_\omega l_\omega}{\sum_{\omega'} \pi_{\omega'} l_{\omega'}} \quad (21)$$

From $\pi_\omega \sim \frac{\rho_\omega}{l_\omega}$ and $\sum_{i=1}^{K} \pi_i = 1$ it also follows that

$$\pi_\omega = \frac{\frac{\rho_\omega}{l_\omega}}{\sum_{\omega'} \frac{\rho_{\omega'}}{l_{\omega'}}} \quad (22)$$

### Spectral analysis of single-molecule footprinting data

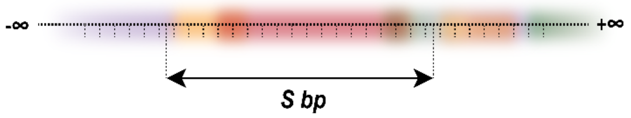


Figure. S9¡. Illustration. for. a. system. used. for. derivation. of probability $P(\gamma,\theta|S)$. of. observing. combination. $\gamma,\theta \in 0,1$. at edges. of. a. randomly. placed. window. with. length. S. bp. on. an infinite. sequence. with. K. footprints. placed. without. positional preferences¡

To explain the frequencies $N^{(\gamma,\theta)}$ of combinations $\gamma,\theta \in 0,1$ observed at different genomic distances from each other in a dataset, and thereby estimate footprint abundancies, we aim to calculate the joint probability $P(\gamma,\theta|S)$ of observing combination $\gamma$ and $\theta$ at both ends of a randomly placed window of size $S$ bp on an infinite sequence (Figure S3) for the stochastic process described above, with footprint prior probabilities $\boldsymbol{\pi} = (\pi_1, \dots, \pi_K)$ and lengths $\boldsymbol{l} = (l_1, \dots, l_K)$, as well as no footprint prior sequence or positional preferences.

Generally, the $P(\gamma,\theta|S)$ can be represented as the sum across all footprints $\omega$,

$$P(\gamma,\theta|S,\boldsymbol{\rho}) = \sum_{\omega} P(\omega) \sum_{i=1}^{l_\omega} P(i|\omega) P(\gamma|i,\omega) P(\theta|S,i,\omega) \tag{23}$$

where $P(\omega)$ and $P(i|\omega)$ are the probabilities that one end hits footprint $\omega$ at position $i$ within this footprint, $P(\gamma|i,\omega)$ is the probability of observing $\gamma$ and $P(\theta|S,i,\omega)$ is probability of observing $\theta$ at another end of the window with length $S$ bp.

Given posterior footprint coverages $\boldsymbol{\rho} = (\rho_1, \dots, \rho_K)$ that can be calculated using (21), and emission probabilities

$$\eta_{\boldsymbol{\omega}}^{(\gamma)} = \begin{cases} \beta^{(\gamma)}, \omega = 1 \\ \alpha^{(\gamma)}, \omega \neq 1 \end{cases} \tag{24}$$

The sum (23) can be written as

$$P(\gamma,\theta|S,\boldsymbol{\rho}) = \sum_{\omega} \eta_{\boldsymbol{\omega}}^{(\gamma)} \frac{\rho_\omega}{l_\omega} \sum_{i=1}^{l_\omega} P(\theta|S,i,\omega) \tag{25}$$

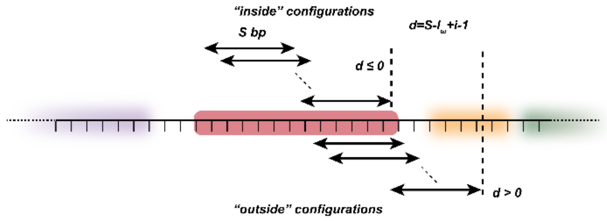


Figure.S0¡.Possible.configurations.of.a.window.overlapping.a. certain.footprint¡

In Figure S4 we can see that $\sum_{i=1}^{l_\omega} P(\theta|S,i,\omega)$ - probability of observing $\theta$ at the right end of the window given that the left end of the window hit a footprint $\omega$, is a sum across two types of configurations $P_{in}(\theta|S,\omega)$ and $P_{out}(\theta|S,\omega)$. Namely, $P_{in}(\theta|S,\omega)$

represents a sum across configurations when both ends are inside footprint $\omega$ ("inside" configurations) and $P_{out}(\theta|S,\omega)$ across configurations when the right end of the window is outside the footprint $\omega$ ("outside" configurations). Therefore,

$$P(\gamma,\theta|S,\boldsymbol{\rho}) = \sum_{\omega} \eta_{\boldsymbol{\omega}}^{(\gamma)} \frac{\rho_\omega}{l_\omega} \sum_{i=1}^{l_\omega} P(\theta|S,i,\omega) = \sum_{\omega} \eta_{\boldsymbol{\omega}}^{(\gamma)} \frac{\rho_\omega}{l_\omega} [P_{in}(\theta|S,\omega) + P_{out}(\theta|S,\omega)] \tag{26}$$

It is obvious that $P_{in}(\theta|S,\omega) = \eta_{\omega}^{(\theta)} N_{in}(\omega,S)$, where $N_{in}(\omega,S)$ represents the number of possible configurations of a window of length $S$ within a footprint $\omega$ and equals

$$N_{in}(\omega,S) = \begin{cases} l_\omega - S + 1, & l_\omega \geq S \\ 0, & l_\omega < S \end{cases} \tag{27}$$

Of note, $N_{in}(bg,S) = 0$ due to minimum window length $S \geq 2$ bp. Therefore, given the footprint emission probability in (24)

$$P_{in}(\theta|\omega,S) = \alpha^{(\theta)} N_{in}(\omega,S), \text{ where } \omega \geq 2 \tag{28}$$

We can represent the sum across "outside" configurations $P_{out}(\theta|S,\omega)$ as a sum across $d = S - l_\omega + i - 1$ - distances from an edge of a footprint $\omega$ to the right edge of a window $S$ (see Figure S4)

$$P_{out}(\theta|\omega,S) = \sum_{d=\max(S-l_\omega,1)}^{S-1} P(\theta|d) \tag{29}$$

where $P(\theta|d)$ represents the probability of observing $\theta$ at distance $d$ from the edge of a footprint. It is obvious that $P(\theta|d)$ can be written as a weighted sum across 2 cases, i.e. if a right edge hit a background or a footprint of any length. Therefore, given the emission probabilities for background and footprints in equation (24) the sum (29) can be represented as

$$
\begin{aligned}
P_{out}(\theta|\omega,S) &= \sum_{d=\max(S-l_\omega,1)}^{S-1} \left(\beta^{(\theta)}P(bg|d) + \alpha^{(\theta)}P(footprint|d)\right) \\
&= \sum_{d=\max(S-l_\omega,1)}^{S-1} \left(\beta^{(\theta)}P(bg|d) + \alpha^{(\theta)}\left(1 - P(bg|d)\right)\right) \\
&= \sum_{d=\max(S-l_\omega,1)}^{S-1} \left(\alpha^{(\theta)} + (\beta^{(\theta)} - \alpha^{(\theta)})P(bg|d)\right) \\
&= \alpha^{(\theta)} \sum_{d=\max(S-l_\omega,1)}^{S-1} 1 + (\beta^{(\theta)} - \alpha^{(\theta)}) \sum_{d=\max(S-l_\omega,1)}^{S-1} P(bg|d) \\
&= \alpha^{(\theta)}N_{out}(\omega,S) + (\beta^{(\theta)} - \alpha^{(\theta)}) \sum_{d=\max(S-l_\omega,1)}^{S-1} P(bg|d)
\end{aligned} \tag{30}
$$

where $N_{out}(\omega,S)$ represents the number of possible "outside" configurations.

When we combine $P_{in}(\theta|S,\omega)$ and $P_{out}(\theta|S,\omega)$ from equations (28) and (30) and plug them into (26)

$$
\begin{aligned}
P(\gamma,\theta|S,\boldsymbol{\rho}) &= \sum_{\omega} \eta_{\boldsymbol{\omega}}^{(\gamma)} \frac{\rho_\omega}{l_\omega}[P_{in}(\theta|S,\omega) + P_{out}(\theta|S,\omega)] \\
&= \alpha^{(\gamma)}\alpha^{(\theta)} \sum_{\omega\geq 2} \frac{\rho_\omega}{l_\omega} N_{in}(\omega,S) \\
&+ \sum_{\omega} \eta_{\boldsymbol{\omega}}^{(\gamma)} \frac{\rho_\omega}{l_\omega}\left[\alpha^{(\theta)}N_{out}(\omega,S) + (\beta^{(\theta)} - \alpha^{(\theta)}) \sum_{d=\max(S-l_\omega,1)}^{S-1} P(bg|d)\right] \\
&= \alpha^{(\gamma)}\alpha^{(\theta)} \sum_{\omega\geq 2} \frac{\rho_\omega}{l_\omega} N_{in}(\omega,S) + \alpha^{(\theta)} \sum_{\omega} \eta_{\boldsymbol{\omega}}^{(\gamma)} \frac{\rho_\omega}{l_\omega} N_{out}(\omega,S) + (\beta^{(\theta)} \\
&- \alpha^{(\theta)}) \sum_{\omega} \eta_{\boldsymbol{\omega}}^{(\gamma)} \frac{\rho_\omega}{l_\omega}\left[\sum_{d=\max(S-l_\omega,1)}^{S-1} P(bg|d)\right] \\
&= \beta^{(\gamma)}\alpha^{(\theta)}\rho_1 N_{out}(\omega = 1,S) + \alpha^{(\gamma)}\alpha^{(\theta)} \sum_{\omega\geq 2} \frac{\rho_\omega}{l_\omega}[N_{in}(\omega,S) + N_{out}(\omega,S)] \\
&+ (\beta^{(\theta)} - \alpha^{(\theta)}) \sum_{\omega} \eta_{\boldsymbol{\omega}}^{(\gamma)} \frac{\rho_\omega}{l_\omega}\left[\sum_{d=\max(S-l_\omega,1)}^{S-1} P(bg|d)\right]
\end{aligned} \tag{31}
$$

Note that $N_{out}(\omega = 1,S) = 1$ as there is only 1 configuration possible when the left edge of a window $S$ is in a background, and $N_{in}(\omega,S) + N_{out}(\omega,S)$ is a total number of possible configurations of a window $S$ overlapping a footprint $\omega \geq 2$ and equals $N_{in}(\omega,S) + N_{out}(\omega,S) = l_\omega$ (see Figure S4). Therefore,

$$P(\gamma,\theta|S,\boldsymbol{\rho}) = \beta^{(\gamma)}\alpha^{(\theta)}\rho_1 + \alpha^{(\gamma)}\alpha^{(\theta)}\sum_{\omega\geq 2}[\rho_\omega] + \left(\beta^{(\theta)} - \alpha^{(\theta)}\right)\sum_{\omega}\eta_{\boldsymbol{\omega}}^{(\gamma)}\frac{\rho_\omega}{l_\omega}\left[\sum_{d=\max(S-l_\omega,1)}^{S-1}P(bg|d)\right]$$
$$= \beta^{(\gamma)}\alpha^{(\theta)}\rho_1 + \alpha^{(\gamma)}\alpha^{(\theta)}(1-\rho_1) + \left(\beta^{(\theta)} - \alpha^{(\theta)}\right)\sum_{\omega}\eta_{\boldsymbol{\omega}}^{(\gamma)}\frac{\rho_\omega}{l_\omega}\left[\sum_{d=\max(S-l_\omega,1)}^{S-1}P(bg|d)\right] \quad (32)$$

Let's consider the sum $\sum_\omega \eta_{\boldsymbol{\omega}}^{(\gamma)}\frac{\rho_\omega}{l_\omega}\left[\sum_{d=\max(S-l_\omega,1)}^{S-1}P(bg|d)\right]$ from the equation (32) more closely. In this sum the $P(bg|d)$ represents probability to find an accessible (background) position at distance $d$ from the right edge of a footprint $\omega$ (see Figure S4). However, as we consider configurations when a certain $\omega$ is fixed, it is clear that this probability is equivalent to the probability to find an accessible position at distance $d$ from an impenetrable barrier in a semi-infinite sequence given the same set of footprints, and, in fact, does not depend on $\omega$ (see Figure S5). Namely, from equations (1) - (4)

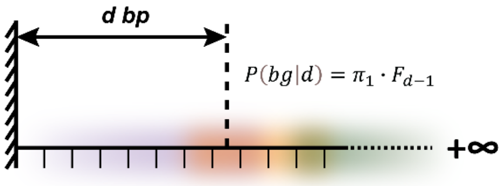

Figure.S❶.Semi_infinite.sequence.with.an.impenetrable barrier¡

$$P(bg|d) = \frac{F_{d-1}\pi_1 B_\infty}{F_\infty} = \pi_1 \cdot F_{d-1} \quad (33)$$

Let $\Omega(s)$ be a cumulative sum $\Omega(S) = \sum_{d=1}^{S}P(bg|d) = \pi_1 \cdot \sum_{d=1}^{S}F_{d-1}$. Then we can split the sum $\sum_\omega \eta_{\boldsymbol{\omega}}^{(\gamma)}\frac{\rho_\omega}{l_\omega}\left[\sum_{d=\max(S-l_\omega,1)}^{S-1}P(bg|d)\right]$ for footprints that are shorter and longer than $S$ and express it in terms of $\Omega(s)$ as follows:

$$
\begin{aligned}
&\sum_{\omega}\eta_{\boldsymbol{\omega}}^{(\gamma)}\frac{\rho_{\omega}}{l_{\omega}}\left[\sum_{d=\max(S-l_{\omega},1)}^{S-1}P(bg|d)\right]\\
&\quad=\sum_{\omega\in l_{\omega}<S}\eta_{\boldsymbol{\omega}}^{(\gamma)}\frac{\rho_{\omega}}{l_{\omega}}\left[\sum_{d=S-l_{\omega}}^{S-1}P(bg|d)\right]+\sum_{\omega\in l_{\omega}\geq S}\eta_{\boldsymbol{\omega}}^{(\gamma)}\frac{\rho_{\omega}}{l_{\omega}}\left[\sum_{d=1}^{S-1}P(bg|d)\right]\\
&\quad=\sum_{\omega\in l_{\omega}<S}\eta_{\boldsymbol{\omega}}^{(\gamma)}\frac{\rho_{\omega}}{l_{\omega}}\left[\sum_{d=1}^{S-1}P(bg|d)-\sum_{d=1}^{S-l_{\omega}-1}P(bg|d)\right]\\
&\quad+\sum_{\omega\in l_{\omega}\geq S}\eta_{\boldsymbol{\omega}}^{(\gamma)}\frac{\rho_{\omega}}{l_{\omega}}\left[\sum_{d=1}^{S-1}P(bg|d)\right]\\
&\quad=\sum_{\omega\in l_{\omega}<S}\eta_{\boldsymbol{\omega}}^{(\gamma)}\frac{\rho_{\omega}}{l_{\omega}}[\Omega(S-1)-\Omega(S-l_{\omega}-1)]+\sum_{\omega\in l_{\omega}\geq S}\eta_{\boldsymbol{\omega}}^{(\gamma)}\frac{\rho_{\omega}}{l_{\omega}}[\Omega(S-1)]\\
&\quad=\Omega(S-1)\sum_{\omega}\eta_{\boldsymbol{\omega}}^{(\gamma)}\frac{\rho_{\omega}}{l_{\omega}}-\sum_{\omega\in l_{\omega}<S}\eta_{\boldsymbol{\omega}}^{(\gamma)}\frac{\rho_{\omega}}{l_{\omega}}[\Omega(S-l_{\omega}-1)]
\end{aligned}
\tag{34}
$$

If we designate $\mathrm{R}\equiv\sum_{\omega'}\frac{\rho_{\omega'}}{l_{\omega'}}$ and therefore $\frac{\rho_{\omega}}{l_{\omega}}=R\cdot\pi_{\omega}$ from equation (22), we can represent the sum (34) as follows

$$
\begin{aligned}
\sum_{\omega}\eta_{\boldsymbol{\omega}}^{(\gamma)}\frac{\rho_{\omega}}{l_{\omega}}\left[\sum_{d=\max(S-l_{\omega},1)}^{S-1}P(bg|d)\right]&=\Omega(S-1)\sum_{\omega}\eta_{\boldsymbol{\omega}}^{(\gamma)}\frac{\rho_{\omega}}{l_{\omega}}-\sum_{\omega\in l_{\omega}<S}\eta_{\boldsymbol{\omega}}^{(\gamma)}\frac{\rho_{\omega}}{l_{\omega}}[\Omega(S-l_{\omega}-1)]\\
&=\Omega(S-1)\cdot R\cdot[\beta^{\gamma}\pi_1+\alpha^{\gamma}(1-\pi_1)]-R\cdot\sum_{\omega\in l_{\omega}<S}\eta_{\boldsymbol{\omega}}^{(\gamma)}\pi_{\omega}[\Omega(S-l_{\omega}-1)]
\end{aligned}
\tag{35}
$$

Finaly, after plugging (35) into (32) and doing simple arithmetic we get

$$
\begin{aligned}
P(\gamma,\theta|S,\boldsymbol{\rho})&=\alpha^{(\gamma)}\alpha^{(\theta)}+\alpha^{(\theta)}(\beta^{(\gamma)}-\alpha^{(\gamma)})\rho_1+R\,(\beta^{(\theta)}-\alpha^{(\theta)})\\
&\cdot\left[\Omega(S-1)\cdot\left(\alpha^{(\gamma)}+\pi_1\big(\beta^{(\gamma)}-\alpha^{(\gamma)}\big)\right)+\sum_{\omega\in l_{\omega}<S}\eta_{\boldsymbol{\omega}}^{(\gamma)}\pi_{\omega}[\Omega(S-l_{\omega}-1)]\right]
\end{aligned}
\tag{36}
$$

Interestingly, when there are no footprints, i.e. $\pi_1=\rho_1=1$ it is easy to show that sum (*36*) reduces to $P(\gamma,\theta|S)=\beta^{(\gamma)}\beta^{(\theta)}$

Also, when the emission probabilities of footprints are the same as emission probabilities of background, i.e. $\alpha^{(\gamma)}=\beta^{(\gamma)}$, which is equivalent to $\alpha^{(\theta)}=\beta^{(\theta)}$, (*36*) reduces to $P(\gamma,\theta|S)=\alpha^{(\gamma)}\alpha^{(\theta)}$, which is constant and it becomes impossible to estimate footprint abundances as background and footprints look identical in the data.

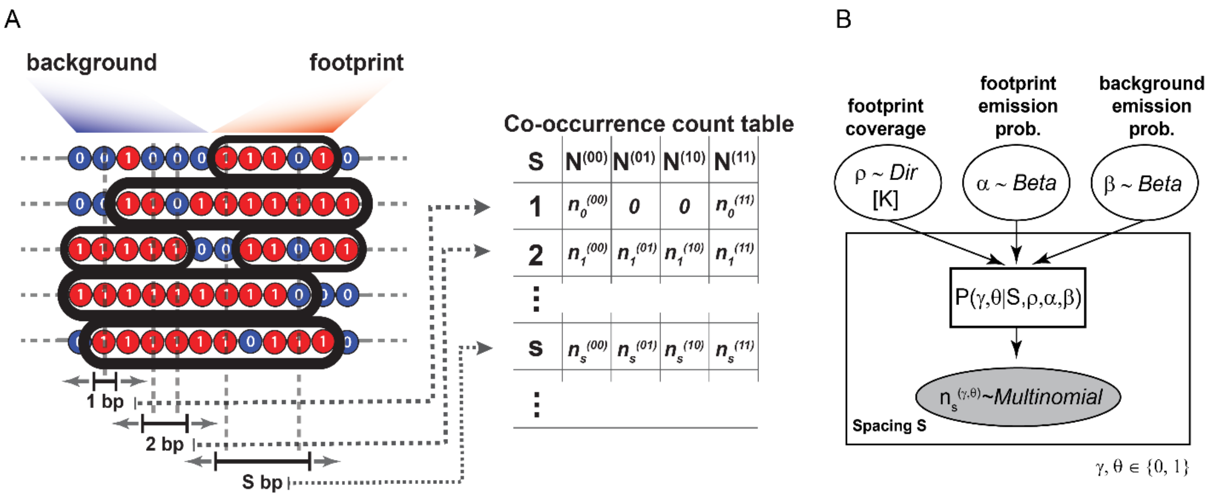


Figure S6. (A) Illustration for creating an observed co_occurrence count table for combinations $\gamma, \theta \in 0{,}1$ at edges of a sliding window with length S bp. (B) Plate notation for the Bayesian model used for inference of footprint abundances and emission probabilities.

Next, for observed co-occurrence counts, as illustrated in the Figure S6A, we can calculate a likelihood function, assuming that counts for each window $S$ follow multinomial distribution and total likelihood function is a product across all available $S$ (Figure S6B). Finaly, assuming Dirichlet prior distribution for footprint coverages $\boldsymbol{\rho}$ and Beta distributions for emission probabilities $\alpha^{(1)}$ and $\beta^{(1)}$, we can construct a Bayesian model and estimate parameters $\boldsymbol{\rho}, \alpha^{(1)}$ and $\beta^{(1)}$ using HMC/NUTS (Hoffman and Gelman, 2014) sampling or Variational Bayes approximation (Kucukelbir et al., 2015) implemented in the probabilistic language Stan (Stan, 2021).

### Predicting footprint positions in single molecule footprinting data

When there are reasonable estimates for footprints lengths, their abundances (either $\boldsymbol{\rho}$ or $\boldsymbol{\pi}$) and emission probabilities ($\alpha^{(1)}$ and $\beta^{(1)}$), we wish to find locations of footprints within each single molecule. In other words, given information about accessible (or methylated) and protected (or unmethylated)

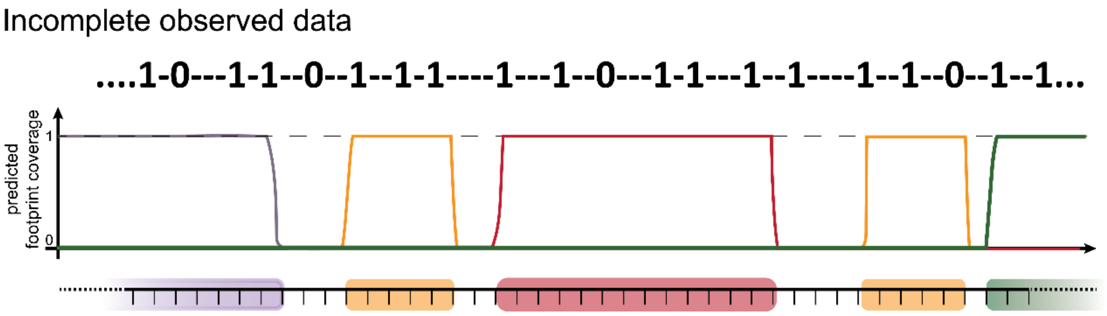


Figure S7. Incomplete data generated by single_molecule footprinting assays and posterior probabilities for each position within a molecule of being covered by each footprint.

positions in a sequenced fragment we aim to recover where footprints of different lengths were located (Figure S7).

To estimate positions for each footprint we can calculate the posterior probabilities that each footprint starts from every position within a sequenced fragment using equation (11) with boundary conditions for infinite sequences (19). To calculate sequence specificity $P(S|n,\omega)$ for a sequence $S$ starting from position $n$ for each footprint $\omega$ we use positional weight matrices of corresponding lengths with entries $\omega_i(S_i \epsilon \{0,1\})$ filled by estimated emission probabilities $\alpha^{(1)} = \alpha$ and $\alpha^{(0)} = 1 - \alpha$ (Figure S8) and $P(S|n,\omega) = \prod_{i=1}^{l_\omega} \omega_i(S_i)$.

| Position within a footprint | Alphabet: 0 | 1 | N(-) |
|---|---|---|---|
| 1 | 1−α | α | 1 |
| 2 | 1−α | α | 1 |
| 3 | 1−α | α | 1 |
| 4 | 1−α | α | 1 |
| 5 | 1−α | α | 1 |
| ⋮ | | | |
| $\ell_\omega$ | 1−α | α | 1 |

Figure.S❹.Positional.weight.matrix.used for.representing.a.footprint.of.length..$l_\omega$ and. used. for. calculating. posterior probabilities¡.

As the resolution of single molecule footprinting assays crucially depends on the density of substrate nucleotides for a chosen enzyme, it is very common that for vast majority of positions within a sequenced fragment information regarding their accessibility is missing and observed data is very sparse (Figure S7). To take this into account we want to calculate $P(S|n,\omega)$ by marginalizing probability across all possible variants of an unobserved part of the sequence $S_{unobs}$ with fixed observed part of the sequence $S_{obs}$, i.e. $P(S|n,\omega) = P(S_{obs}|n,\omega) \cdot \sum_{S_{unobs}} P(S_{unobs}|n,\omega)$. It is easy to show that $\sum_{S_{unobs}} P(S_{unobs}|n,\omega) = 1$ and $P(S|n,\omega) = P(S_{obs}|n,\omega)$. Indeed, at any positon with unobserved data there is only 2 variants possible, i.e. with 0 or 1 at this position, and as we assume independence of positions within PWM and the sum across these 2 variants is always 1, the sum $\sum_{S_{unobs}} P(S_{unobs}|n,\omega)$ equals 1. Therefore, we can introduce a letter in the alphabet which will correspond to missing data (NA or (–) as in Figure S7) and add an extra column into a PWM for this letter filled with ones (Figure S8).
